# Landscape genomics highlights the adaptive evolution of chickpea across the Silks Roads

**DOI:** 10.1101/2024.06.06.597750

**Authors:** Lorenzo Rocchetti, Giulia Frascarelli, Monica Rodriguez, Alice Pieri, Simone Papalini, Luca De Antoni, Erika Vitali, Astrid Vincze, Roberto Biello, Andrea Benazzo, Creola Brezeanu, Chiara Santamarina, Elisa Bellucci, Laura Nanni, Marzia Rossato, Massimo Delledonne, Elena Bitocchi, Roberto Papa

**Author notes:** Author for correspondence: Roberto Papa.

## Abstract

Environmental heterogeneity and human-mediated dispersal have jointly shaped the genetic diversity and local adaptation of crop species. Understanding the genetic basis of these processes is essential for improving crops across diverse agro-environmental conditions.

We characterized population structure and geographic patterns of genetic diversity in 532 chickpea genotypes spanning most of the cultivated range. Using redundancy analysis on 208 georeferenced landrace-derived genotypes from the Mediterranean Basin to Central Asia, we identified genotype-environment associations (GEAs) and traced adaptive variation along historical Silk Road routes.

Both environmental and geographic factors significantly shaped chickpea diversity. GEA loci were enriched for genes involved in heat and drought tolerance, while key geographic regions harbored reservoirs of adaptive alleles and early-flowering genotypes, supporting flowering time as an escape strategy from terminal stress.

These findings provide a genomic framework for integrating climate-adaptive alleles into chickpea breeding programs targeting drought- and heat-prone environments.

## Introduction

Plant-based diets, and particularly the widespread utilization of legume-based foods, can improve human health, support biodiversity conservation, and mitigate the environmental impact of modern agriculture, including its contribution to climate change, while meeting the nutritional demands of a growing global population ^1–3^. In this context, increasing the cultivation and consumption of grain legumes represents a valid strategy to facilitate the adoption of healthy diets, ensure a sufficient intake of plant-based protein while respecting the planetary boundaries by reducing agronomic inputs such as biological nitrogen fertilization^4^. The conservation and characterization of food legume genetic resources are therefore paramount, as they provide the foundation for developing new varieties that can improve food legume production and promote their consumption ^4^.

The levels of genetic diversity in cultivated species are largely determined by the diversity present in their wild progenitors ^5^. However, an extensive agronomic and morphological diversity is observed in crops as a result of their dispersal and adaptation to different agro-environments out of the center of domestication ^6–10^. Understanding the structure of crop diversity in relation to spatial and ecological variation is crucial to elucidate the role of different evolutionary forces during adaptation. Landscape genomics has emerged as a powerful framework to investigate patterns of adaptive genetic variation across environmental gradients. Initially developed for ecological studies^11–14^, it has increasingly been applied to crop systems to assess adaptive potential and local environmental responses ^15,16^. By detecting genotype–environment associations (GEAs), landscape genomic analyses can help to elucidate the genetic architecture of local adaptation and ultimately infer the adaptive potential of a given genotype or population under contrasting environmental conditions. Although landscape genomics approaches have recently been applied to crops, their use in legume species remains limited. Notably, landscape genomics analyses have been conducted on American wild and domesticated common bean germplasm to investigate the effects of demographic processes and local selection on genetic variation ^17^, as well as to examine the influence of climatic adaptation on mung bean dispersal routes across Asia ^18^.

Chickpea (*Cicer arietinum* L.) is the second most important food legume, after the common bean, for direct human consumption^19^. It was domesticated in Southern Eastern Turkey ^20,21^ and subsequently spread along the Silk Roads to the diverse agro-environments of the Mediterranean Basin, North Africa, East Africa, Central Asia, and the Indian subcontinent. Today, it is grown as a spring and summer crop in the Mediterranean Basin, North America, and Australia, and as a post-monsoon winter crop in Ethiopia and India. Two major chickpea market classes are distinguished: desi, characterized by purple flowers and small dark seeds, predominantly grown in the Indian sub-continent and Ethiopia, and kabuli, characterized by white flowers and larger, light-colored seeds with a thin seed coat, mainly grown and consumed in the Mediterranean basin, the Middle East, and North America.

Several studies have used large collections of chickpea genetic resources to study patterns of diversification based on biological status (e.g., wild *vs* domesticated) or seed shape type classes (desi *vs* kabuli) ^22,23^. However, the effect of spatial and environmental factors on the population differentiation has only been addressed in a local study conducted on 77 Pakistani landraces grown at different elevations along an isothermal gradient^24^.

Here, we assembled a large collection of single seeds descent lines ^25,26^ covering the major areas of chickpea cultivation. We analyzed the resulting sequence data using population and landscape genomics, providing new insights into the post-domestication spread of this crop. Additionally, using a subset of 208 geo-referenced genotypes, derived from landraces populations, spanning from the Mediterranean basin to Central Asia, we examined the relative effects of geography, environment, and population structure on genetic diversity. Ultimately, we searched for candidate genomic regions associated with major environmental variables and mapped their spatial variation across the Silk Roads.

## RESULTS and DISCUSSION

### Population structure analysis of chickpea germplasm reveals a geographic pattern of diversification

Following domestication in Middle East, chickpea spread from its area of origin and adapted to diverse agroclimatic zones ^25^ leading to wide phenotypic and genetic variability that can be observed in domesticated chickpea genetic resources ^23,26–31^. Deciphering the evolutionary pathways that enable the crop to adapt in diverse agro-environment, present a valuable opportunity to enhance plant breeding toward adaptative traits and consequently to understand the molecular basis of adaptation ^8^. We therefore investigated the genetic structure of chickpea germplasm according to geography and agricultural environment.

We assembled a panel of 532 chickpea lines obtained by single-seed descent (SSD) from diverse accessions^25,26^. Unlike previously assessed collections, our panel included a substantially larger proportion of Mediterranean and European genotypes^23,28^, providing broader geographic coverage of an important area of the crop’s post-domestication range.

Admixture analysis did not identify a clear number of population subgroups. The cross-validation coefficient decreased exponentially from K = 2 to K = 8 without reaching a minimum (**Fig. S1**), consistent with a pattern of continuous genetic differentiation rather than discrete clustering, as also reported in previous studies^23,28,30,32^. This level of introgression likely reflects the dissemination of the crop in response to political factors and complex trade routes, analogous to patterns observed in other crops such as common bean ^9^. We therefore examined meaningful geographic subdivisions alongside *desi* and *kabuli* differentiation across subsequent K values (**Fig. 1; Fig S2**). Considering the crop expansion out of its center of origin in Southern Turkey, we divided the collection into 13 representative macro-geographic area based on each genotype’s country of origin (**Fig. 1**).

**Fig. 1:**
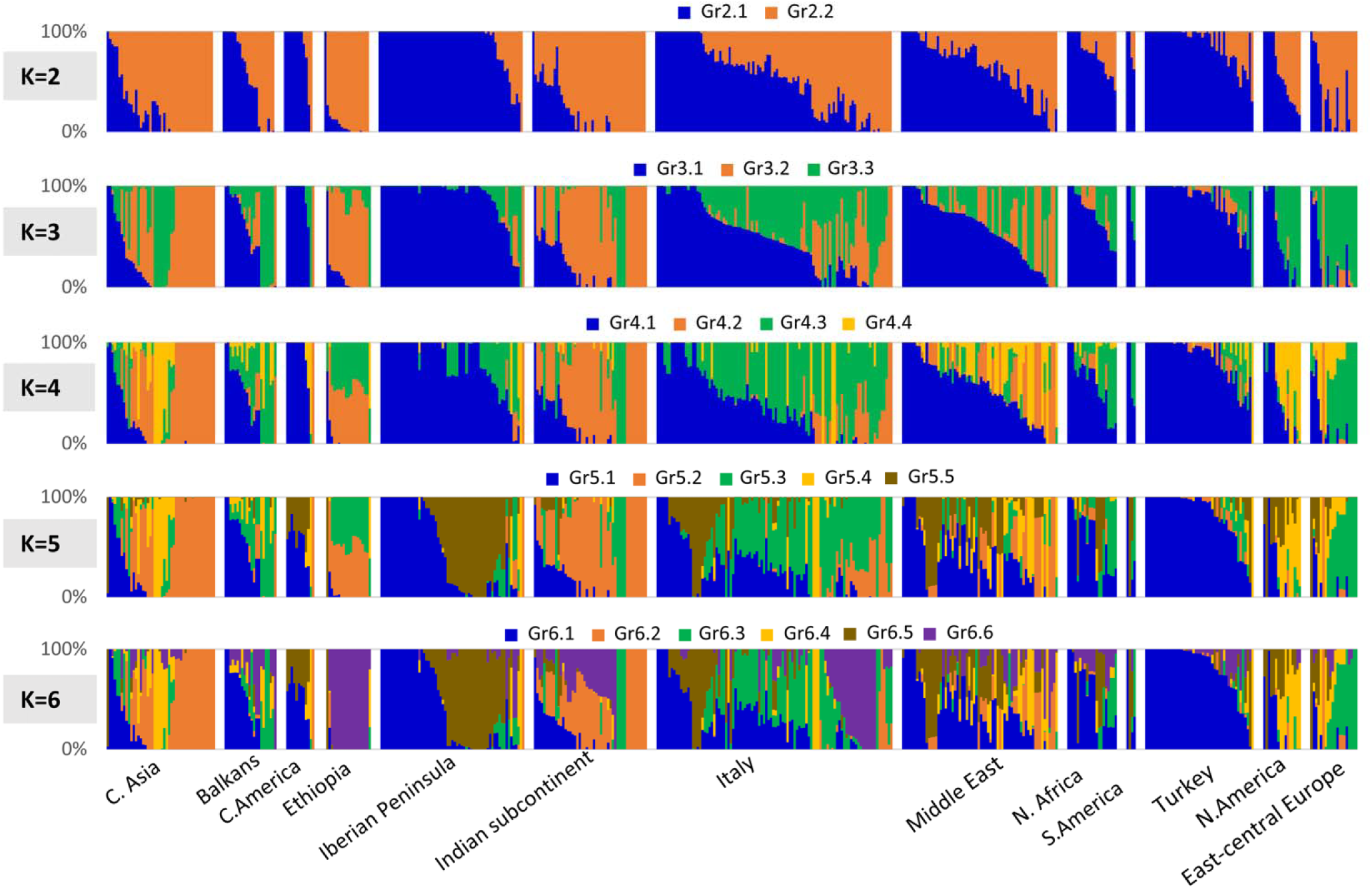
Population structure analysis performed with ADMIXTURE based on 2,050 SNPs across 532 chickpea accessions. Individuals are grouped following the defined 13 major geographic areas.

At K=2, two groups were identified (Gr2.1 and Gr2.2) separating genotypes from the Mediterranean basin (average *q*_Gr2.1_ > 0.85) from those of Central Asia, the Indian subcontinent, Ethiopia and Eastern Europe (average *q*_Gr2.2_ > 0.82) (**Fig. 1**; **Fig. S2**). A more balanced representation of the two groups was present in Italy, the Balkans and the Middle East (**Fig. 1**), with a high proportion of admixed genotypes (**Fig. S2**). At K=3, Gr3.1 corresponded to Gr2.1 identified at K=2, whereas Gr3.2 and Gr3.3 represented subdivisions of Gr2.2. Gr3.2 mainly included genotypes from the Indian subcontinent, Central Asia, and Ethiopia with high Gr2.2 membership, whereas Gr3.3 comprised mostly admixed Gr2.2 genotypes and a subset with high *q*_Gr2.2_, primarily from Europe, the Caucasus, and the USA (**Fig. 1**; **Fig. S2**). At K=4 (**Fig. 1**), Gr4.1 and Gr4.2 corresponded to Gr3.1 and Gr3.2 at K=3, whereas Gr4.3 and Gr4.4 represented subdivision of Gr3.3 at K=3. Notably, Gr4.4 included genotypes from Eastern Europe, Caucasus, the Middle East and Central Asia (**Fig. 1**). At K=5, we observed a differentiation between the western and eastern sides of the Mediterranean basin, particularly between the Iberian Peninsula and Turkey. Additionally, a meaningful subdivision occurred at K=6, where the new group Gr6.6 clearly differentiated the entire Ethiopian germplasm. The same ancestry is largely present in Italy and the Indian continent but absent in Central Asia.

To examine the geographic distribution of genetic structure, we used the 208 geo-referenced genotypes covering the Mediterranean Basin and West and Central Asia, spatially interpolating membership coefficients (q*_i_* > 0.4) for both K=3 and K=6 (**Fig. 2**). K=3 was selected because it represents an inflection point in the cross-validation (CV) error decay curve, while K=6 was selected because it corresponds to the partitioning level at which the CV error begins to plateau and where the whole set of Ethiopian genotypes form a distinct group (**Fig. S1**). We confirmed on both partitioning the same groups in the Anatolian region and Central Asia specifically highlighted in blue and orange respectively. At K=6 we confirmed a further differentiation between eastern and western sides with the group Gr6.1 mainly present in Turkey with signatures in North Africa and the group Gr6.5 strongly present in the Iberian Peninsula. The central part of the Mediterranean basin represented by Italy resulted highly admixed with several genetic groups including Gr6.3, Gr6.5 and Gr6.6. (**Fig. 2**).

**Fig. 2.**
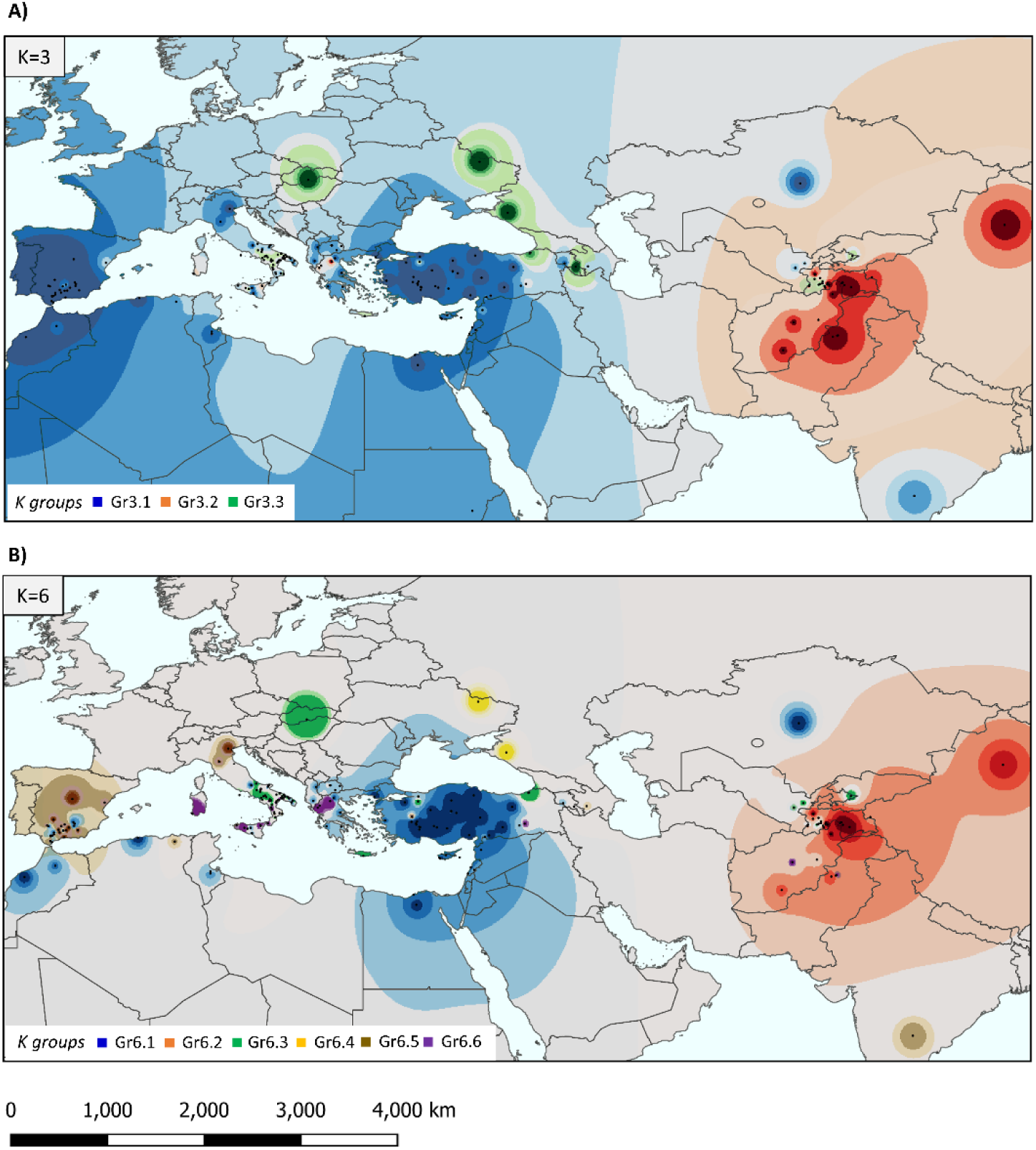
Spatial distribution (Inverse Distance Weighted, IDW) of the interpolated membership coefficient q*_i_>0.4* from *Admixture* analysis at K=3 and K=6. Colors indicate ADMIXTURE group assignments at both K=3 and K=6 partition.

Considering that most of research and breeding programs distinguish chickpeas based on their market types desi and kabuli, we compared the population structure results of K=2 and K=3 according to this dichotomy. Using a membership coefficient threshold of q*_i_* > 0.90, at K=2 we defined 164 and 198 individuals as pure Gr2.1 and Gr2.2 genotypes, respectively. Both market types were present in both groups, but differed significantly in frequency (χ² test, P < 0.001; **Fig. S3a**). Gr2.1 featured a higher proportion of kabuli than Gr2.2 (69% vs. 13%), while the opposite was true for desi (31% vs. 87%). These results agree with previous studies ^23,32^ where Western and Eastern areas subdivision also reflected a significant differentiation between the two crop market types, with western regions featuring more kabuli types and Central Asia and the Indian subcontinent featuring mostly desi types (**Fig. S4b**). Significant differences between the q*_i_* of desi and kabuli were also found among the entire set of 532 genotypes (**Fig. S3b**).

At K=3, we assigned 142, 53 and 45 pure individuals to Gr3.1, Gr3.2 and Gr3.3 (q*_i_* > 0.90). An unrooted phylogenetic tree constructed from these pure genotypes revealed a meaningful differentiation between desi and kabuli types. Specifically, we found both desi and kabuli genotypes in Gr3.1 and Gr3.3, while kabuli types were absent in Gr3.2, with the only exception being an Italian kabuli genotype (**Fig. 3A**). The phylogenetic tree also displayed a clear geographic structure (**Fig. 3A**). Gr3.1 comprised four clusters: one predominantly consisting of Turkish and Middle Eastern genotypes, two featuring Iberian Peninsula genotypes, and one composed of Italian and North African genotypes. Gr3.2 was characterized by three clusters separating the Central Asian, Indian subcontinent and Ethiopian genotypes. Gr3.3 comprised one cluster representing Eastern European germplasm and one including Central Asian genotypes alongside breeding materials from the USA and European cultivars. Taken together, these results support the evidence that desi was the early domesticated type, which spread from the area of domestication, whereas kabuli traits arose multiple times following desi diffusion, mainly in the West Asia, Mediterranean basin and Central Asia^20,28^.

**Fig. 3.**
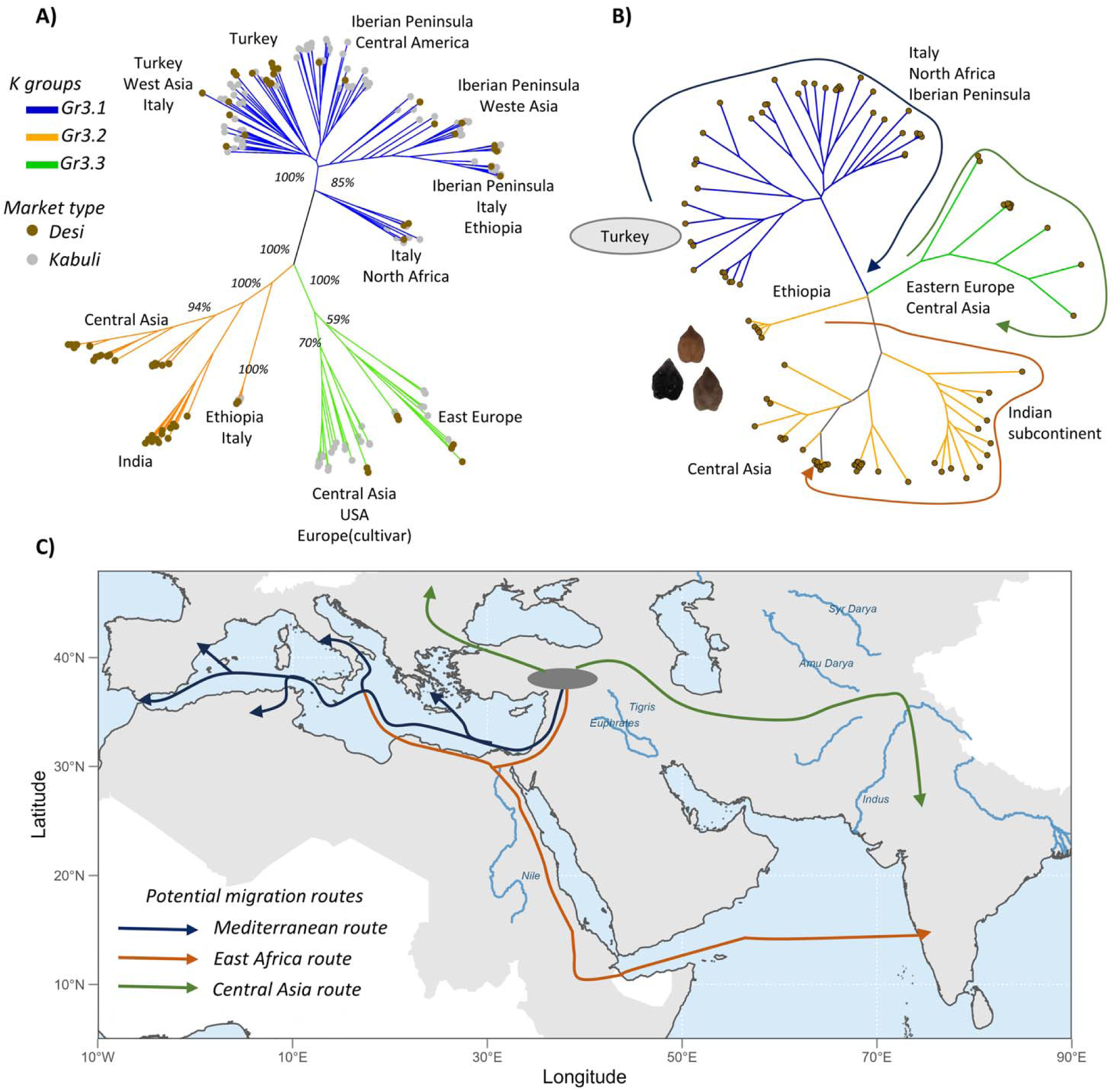
Neighbor-joining (NJ) phylogenetic trees and candidate migration routes **(a)** NJ tree based on 240 individuals with q*_i_* > 0.90 for Gr3.1, Gr3.2 and Gr3.3. Branch colors indicate membership of Gr3.1 (blue), Gr3.2 (orange) and Gr3.3 (green). Brown and gray circles indicate desi and kabuli types, respectively. Bootstrap values ≥50% are shown for the major nodes. **(b)** NJ tree based on 101 desi genotypes with q*_i_* > 0.90 for Gr3.1, Gr3.2 and Gr3.3. Geographic labels are reported for each branch, with the center of domestication (Turkey) highlighted in dark grey. **(c)** Potential migration routes of chickpea along the Silk roads.

To obtain a clearer picture of chickpea migration outside the domestication center, we constructed a phylogenetic tree using only minimally introgressed desi genotypes, thereby removing the confounding signal of later-differentiated kabuli types.(**Fig. 3B**). The desi-only tree supports the same separation into three genetic groups. Notably, among Gr3.2 individuals, Ethiopian genotypes were more closely related to Gr3.1, which predominantly comprised Turkish and Middle Eastern genotypes, than to the remainder of Gr3.2. (**Fig. 3B**). These results support one of the main reported pathways of chickpea diffusion through the Mediterranean, as shown by the strong relationship between Turkish and European (mainly Italy and Iberian Peninsula) and North African accessions ^29^. At the same time, they are consistent with more recent studies highlighting a complex dispersal scenario for chickpea in Ethiopia and Asia ^32^. Specifically, the intermediate position of Ethiopian chickpea between Turkish/Mediterranean and Asian clusters in the phylogenetic tree does not support the currently suggested migration of chickpea into India and subsequently toward Ethiopia ^29^.

We propose instead that Mediterranean chickpea reached Ethiopia via an East African route, moving across the Nile River corridor or through the marine trade network across the Red Sea, before subsequently spreading to India across the Indian Ocean (**Fig. 3C**). This dispersal route, connecting Roman-Egyptian ports to the Indian subcontinent, is described in detail in the Periplus of the Erythraean Sea, an ancient Greco-Roman navigational text that documents the commercial maritime networks of the first century CE. Support for Abyssinia as a connecting point between the Mediterranean and South Asia is further supported by the population structure detected at K=6, where Ethiopian genetic ancestry is largely present in the Italian and Indian germplasm, but not in Central Asian genotypes. A comparable dispersal pattern has also been proposed for sorghum ^33^, suggesting that the *guinea* race moved from Western to Eastern Africa and then reached India by sea trade across the Indian Ocean.

PCA analyses also underscored the strong genetic affinity between Mediterranean and Ethiopian accessions (**Fig. S4a**). PC1 and PC2 explained 14% and 8% of the total genetic variation, respectively. The score plot confirmed a geographical pattern, with PC1 separating Turkish germplasm from Central Asia and the Indian subcontinent. Mediterranean genotypes were broadly distributed but clustered mainly near the Turkish group. Consistent with admixture results, PC1 separated Gr2.1 and Gr2.2 at K=2, while at K=3 PC1 separated Gr3.1 from Gr3.2 and PC2 distinguished Gr3.3 from Gr3.2 (**Fig. S4c**). Desi and kabuli types overlapped in the bottom-right of the plot; however, the bottom-left was dominated by desi genotypes, whereas genotypes with increasing PC2 values were predominantly kabuli (**Fig. S4b**).

Collectively, population structure, NJ tree, and PCA analyses robustly revealed the close genetic relationship between Mediterranean accessions and Ethiopian germplasm ^32^. Archeological records suggest that chickpea spread from the Middle East to South Asia at least 4000 years ago and reached Ethiopia at least 2300 years ago^34^. Interestingly, the records from Northern India are more ancient (Late Neolithic, 5450−3500 BC) than those from Southern India (Bronze Age, 2800−1300 BC; Iron Age, 1300−500 BC; Recent, 300 BC to 200 AD), This temporal gradient may reflect two distinct introduction events: Northern Indian genotypes likely originated from Turkey and dispersed eastward via the Middle East, whereas the more recent archaeological signal in Southern India may reflect a later introduction of Ethiopian chickpea through Roman-era Eastern trade networks, following the Roman conquest of Egypt by Augustus in 30 BC and the subsequent intensification of maritime trade across the Red Sea and Indian Ocean ^34^.

### Signatures of environmental adaptation

Within the full collection of 532 accessions, we identified 208 geo-referenced accessions, of which 202 are landraces, primarily distributed across the Mediterranean Basin and extending into West and Central Asia. As landraces of an annual crop, these accessions very likely represent locally adapted material, providing a valuable opportunity to investigate how landscape variation, encompassing both geographic and environmental factors, shaped genetic diversity along the Silk Roads. We first assessed the influence of geographic distance on genetic variation using a Mantel test with 1,000 permutations to evaluate the correlation between geographic and genetic distance matrices. A significant positive correlation was detected (r = 0.34, P < 0.001), indicating that isolation by distance (IBD) has strongly influenced the spatial structuring of genetic diversity across the collection (**Fig. S5**). To disentangle the relative contributions of environmental, geographic, and demographic factors to genetic variation, we applied redundancy analysis (RDA) as a multivariate framework enabling simultaneous correlation of the genetic diversity matrix with a landscape matrix comprising environmental variables, geographic variables, and population structure covariates. This approach allows to dissect the overall influence of landscape variation of genetic diversity as well as the single contribution of environment, geography and genetic structure, enabling the identification of genomic variation associated with the joint action of multiple climatic variables. The RDA full model revealed that 34% of total genetic variance was explained by the combination of population structure, geography, and environmental variables. We then used partial RDA (pRDA) to estimate the variance proportion explained by one set of explanatory variables when the effect of the others has been removed by considering them as covariates. The pure environmental effect was significant and explained 3% and 8% of the total genetic variance and constrained variance, respectively (**Table 1**). Population structure was also significant and had the strongest effect, it explained the 22% of the total genetic variance (64% of the variance explained by the full model), highlighting the role of demographic history in shaping chickpea genetic variation. The geographic effect was also significant and explained 2% of the total genetic variance (5% of the explained variance) (**Table 1**).

**Table 1.** Partial redundancy analysis, showing the influence of environment (bio8 + bio9 + bio15 + bio18 + bio19), geography (latitude + longitude), and population structure (PC1 + PC2) on genetic variation. The proportion of explained variance represents the total constrained variation explained by the full model.

| RDA model | Variance | Proportion of explained variance | Proportion of total variance | Pr>F |
| --- | --- | --- | --- | --- |
| Full model: $Y \sim \text{Env} + \text{Geo} + \text{Struct}$ | 12,590 | 1 | 0.34 | *** |
| Pure environment: $Y \sim \text{Env} \mid (\text{Geo} + \text{Struct}.)$ | 9,837 | 0.08 | 0.03 | *** |
| Pure structure: $Y \sim \text{Struct} \mid (\text{Env} + \text{Geo}.)$ | 8,002 | 0.64 | 0.22 | *** |
| Pure geography: $Y \sim \text{Geo} \mid (\text{Env} + \text{Struct}.)$ | 661 | 0.05 | 0.02 | *** |
| Confounded Env/Geo/Struct | 2,943 |  | 0.08 |  |
| Environment and geography: $Y \sim \text{Geo} + \text{Env}.$ | 4384 | 0.34 | 0.12 | *** |
| Unexplained | 23,460 |  | 0.65 |  |
| Total | 36,050 |  |  |  |

To visualize the role of environmental and geographic factors on genetic group divergence, we applied an RDA model where the selected environmental variables and geographic coordinates were used as explanatory components of genetic diversity. The model was significant (P<0.05) and explained 12% of the total genetic variance (**Table 1**). The RDA biplot derived from this model showed that 58% and 19% of the total constrained variation is explained by the first and second RDA components, respectively (**Fig. 4**). By highlighting genotypes according to specific genetic groups (q*_i_* > 0.9) for the two partitioning at K=3 and K=6, we identified interesting relationships between genetic groups across environmental and geographic variables. The groups Gr3.1 and Gr3.3 were differentiated from the others based on temperature in the wettest quarter (bio8) and precipitation in the warmest quarter (bio18). The same difference occurs between the groups Gr6.1 and Gr6.2. The subdivision between the group Gr6.1 and Gr6.5 along the RDA2 component is associated to differences for the variable bio8 and bio18. The orange group largely present in Central Asia represented by Gr3.2 and Gr6.2 was distinguished from all the others based on longitude, high temperature in the driest quarter (bio9) and low winter precipitation (bio19) (**Fig. 4**). This indicates that this group could provide an important source of drought-adaptive alleles. As already observed in soybean, environmental factors, such as temperature and latitude, play a significant role in driving the divergence of wild population subgroups^35^.

**Fig. 4.**
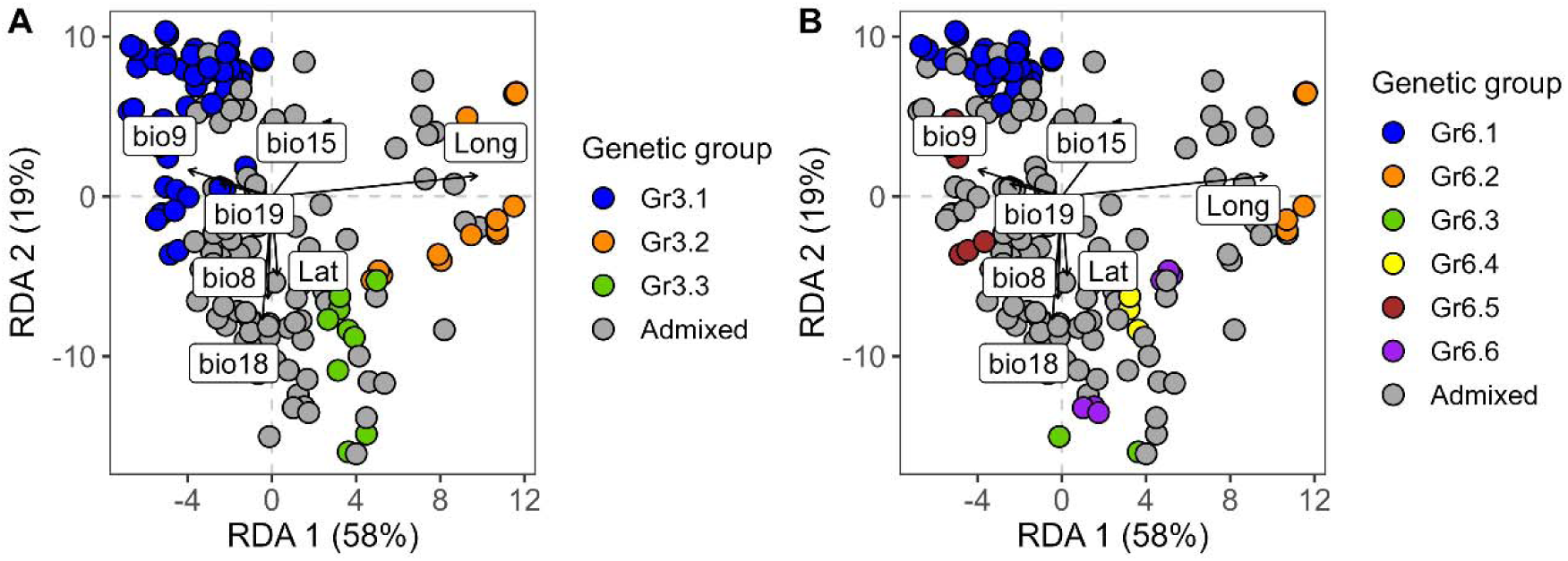
Redundancy analysis biplot relating genotypes (circle) with environmental (bio8, bio9, bio15, bio18, bio19) and geographic (latitude and longitude) variables (arrows). Genotypes with high ancestry coefficient (*qi* values >0.90) are colored highlighting the specific genetic groups at a) K=3 and b) K=6

### Genotype–environment association

By applying RDA, we identify genomic variants associated with environmental variation (GEA-QTLs) in chickpea, representing a major step toward integrating genotype–environment associations into legume crop genomics. Specifically, we used partial RDA (pRDA) to identify covarying loci in response to the multivariate environments derived from temperature (bio8 and bio9) and precipitation (bio15, bio18 and bio19) variables (**Fig. 5**). Geographic coordinates and genetic structure variation were included as covariates to account for their confounding effects. We acknowledge that expanding this approach to include additional environmental factors, such as soil properties, could further refine the characterization of agro-environmental adaptation and uncover novel associations.

**Fig. 5.**
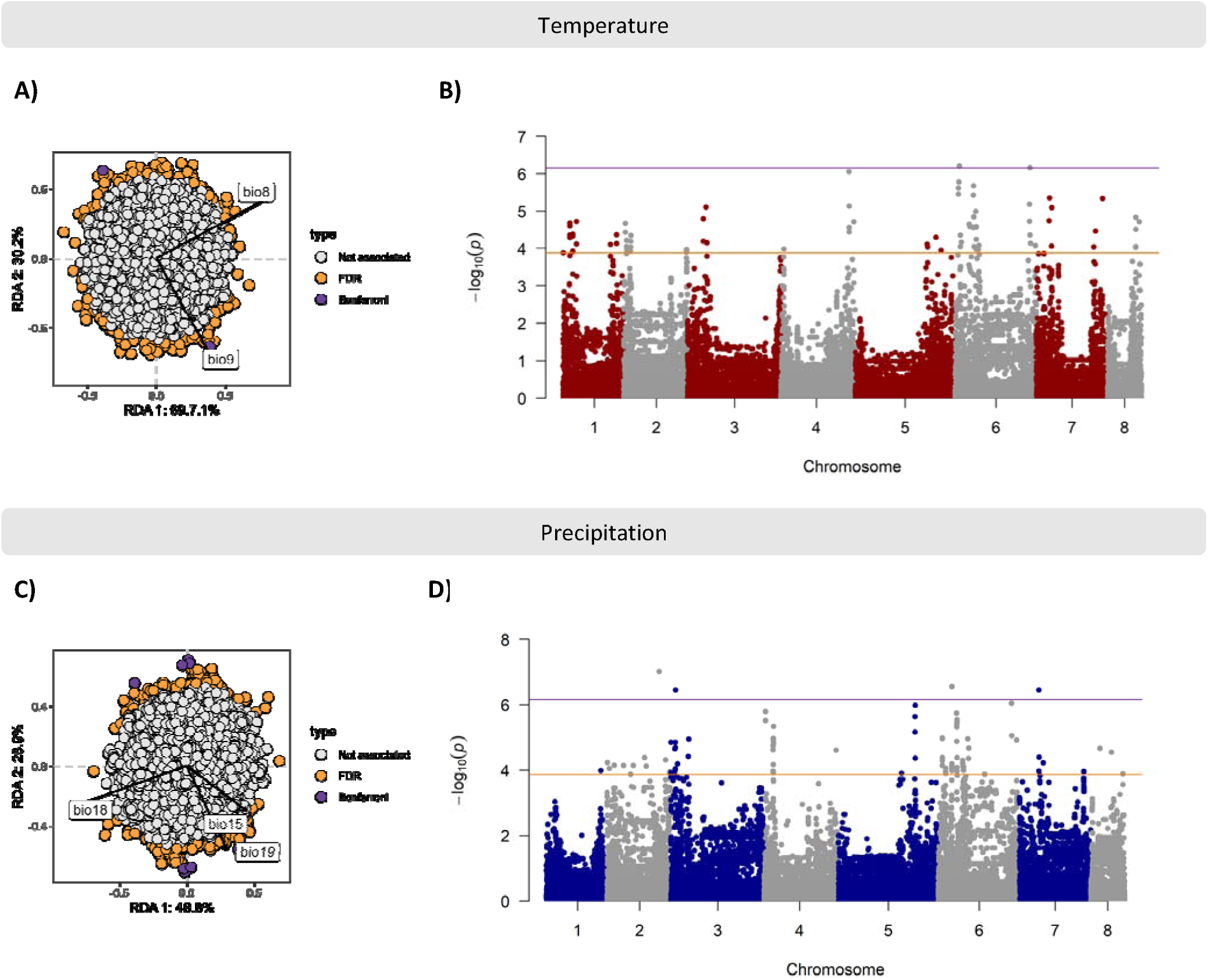
Genotype–environment associations for temperature variables bio8 and bio9 (**a, b**), and precipitation variables bio15, bio18 and bio19 (**c,d**), using partial RDA. **(a, c)** Projection of SNP alleles and environmental variables. Outliers are identified by extremeness along a distribution of Mahalanobis distances estimated between each locus and the center of RDA space (RDA1, RDA2). **(b, d)** Manhattan plot with the distribution of -log_10_(p values). Two threshold levels were applied: Bonferroni P<0.05/number of comparisons (purple line) and Storey FDR (q-value; q < 0.05) (orange line).

Considering temperature variables, the pRDA was significant (P = 0.002) and accounted for 1% of total variance. The first RDA axis accounted for 69.7% of the constrained variance and was associated with bio8 (mean temperature in the wettest quarter), whereas the second axis explained 30.2% of the constrained variance and was correlated with bio9 (mean temperature in the driest quarter). The multivariate GEA method identified 94 loci using a Storey false discovery rate (FDR) (q-value; q < 0.05) correction; among them one locus, on chromosome 6 showed significant associations using a Bonferroni correction (P<0.05 **Fig. 5B, Fig. S6, Table S3**).

For precipitation variables, the pRDA was significant (P=0.006) and accounted for 1.5% of total variance. The first RDA axis explained 48.8% of the constrained variance and was associated with bio18 (amount of precipitation in the warmest quarter), whereas the second axis accounted for 28.9% of the constrained variance and was mainly correlated with bio19 (amount of precipitation in the coldest quarter) (**Fig. 5C**). Overall, 97 loci were significantly associated with the combination of the three variables (bio15, bio18 and bio19) using a Storey FDR (q-value; q < 0.05) correction. However, after applying the Bonferroni correction, the number of strongly associated loci was reduced to four, located on chromosomes 2, 3, 6, and 7 (**Fig. 5D**, **Fig. S6, Table S3**).

The same associations for temperature and precipitation were performed correcting for population structure using five principal components (PCs). For both temperature and precipitation associations, we found significant correlations between the respective logarithm of P-values, with explained variances greater than 70% (R^2^>0.7) (**Fig. S7, Fig. S8**). An additional GEA analysis was performed using envGWAS ^36^, which is based on a univariate approach. No significant loci were associated with the bioclimatic variables already studied with the RDA (i.e., the temperature variables bio8 and bio9, and the precipitation variables bio15, bio18, and bio19), apart from 7 markers on chromosomes 2, 3, 5 and 7 that were significantly associated with bio9 (**Fig. S9**). In contrast to the RDA results, no markers located on chromosome 6 were associated with any temperature variables. The discrepancy between the two GEA approaches tested in this work may be explained by the fact that envGWAS analyzes a single bioclimatic variable and a single locus at a time. Consequently, it lacks the statistical power to capture the genetic architecture of complex polygenic traits, such as responses to temperature and drought ^37^. The significant GEA loci identified using a Bonferroni correction threshold were queried across the full panel of 532 genotypes, with the ultimate aim of providing users of our collection with a practical tool to identify genotypes carrying potentially beneficial alleles for drought and heat stress tolerance (**Table S1**).

To identify candidate genes, we performed linkage disequilibrium (LD) analysis using PopLDdecay. The results indicated extensive genome-wide LD, with r^2^ decaying to 0.3 at approximately 500 kb, but exhibiting a slower decay thereafter, reaching r^2^ = 0.2 at approximately 2000 kb (**Fig. S9**). While these values are comparable to other self-pollinated legumes such as common bean ^9^ they are larger than the previous reported values for chickpea ^29,38^.

We hypothesize that this difference is primarily due to the use of a more contiguous reference genome assembly (∼529 Mb) compared to the previous version (∼347 Mb; ^39^) used in earlier studies. This increased contiguity captures larger physical intervals between recombination events that were likely underestimated in more fragmented assemblies. Furthermore, our panel likely possesses a distinct genetic structure compared to the global set used other works ^29^, due to a higher representation of Mediterranean genotypes that contrast with Asian and Indian germplasm. Therefore, the observed LD extent likely reflects differences in genome assembly scaling and panel composition rather than a biological departure from known recombination patterns in chickpea.

To remain conservative and avoid overestimating candidate regions, we constrained our search to a 100 kb window flanking each associated marker, following the high-stringency approach already applied in common bean ^9^. This allowed us to prioritize genes in the closest physical proximity to the peak association signals.

We found 54 genes in the 100-kbp window surrounding the five SNP markers associated with temperature and precipitation variables (Bonferroni P<0.05/number threshold) (**Note S1 and Table S3**, including related positions in the CDC Frontier v1.1 reference genome). A detailed survey of gene functions was carried out, mainly using *Arabidopsis* orthologs (**Note S1**). Several candidate genes (17 of 54) were found to be involved in responses to abiotic stress factors such as heat, drought, and salinity. GO analysis conducted against the *Arabidopsis* genome highlighted significant enrichment in the biological processes related to biotic and abiotic stress responses as well as terms related to cell membrane components (**Fig. S11**). These GO terms have been already identified as overexpressed in response to both biotic and abiotic stresses in potato ^40^ and rice ^41^. Excluding genes of unknown function, we found that almost all remaining genes were associated with mechanisms used by plants to survive and adapt to environmental changes, such as abscisic acid (ABA), gibberellin (GA), cytokinin (CK), salicylic acid (SA), jasmonic acid (JA) and auxin signaling, cell wall and cuticle biosynthesis, growth and leaf senescence. The most promising candidate genes are listed in **Table 2**. As example, we cite five closely linked homologous genes (Ca2g201700, Ca2g201800, Ca2g201900, Ca2g202000 and Ca2g202100) coding for U-box domain-containing protein 21-like. The orthologous gene in *Arabidopsis*, AT5G37490 gene, encodes for a plant U-box type E3 ubiquitin ligase (AtPUB21, AtCMPG5). Comparative analyses of *Arabidopsis* gene expression under low, optimal, and high growth temperatures have clarified how root growth adapts to elevated ambient temperatures and identified AtPUB21 as one of the genes strongly downregulated in response to heat stress ^42^. Additionally, transcriptomic analyses of leaves from wild-type creeping bentgrass (Agrostis stolonifera cv. “Penncross”) and an isopentenyl transferase (ipt)–overexpressing line (S41), which exhibits enhanced cytokinin levels and drought tolerance, revealed that the AtPUB21 ortholog was among the downstream genes differentially expressed under drought stress. These findings suggested that improved drought tolerance observed in ipt-overexpressing plants was associated with ubiquitin-mediated proteolysis pathways ^43^.

**Table 2.**
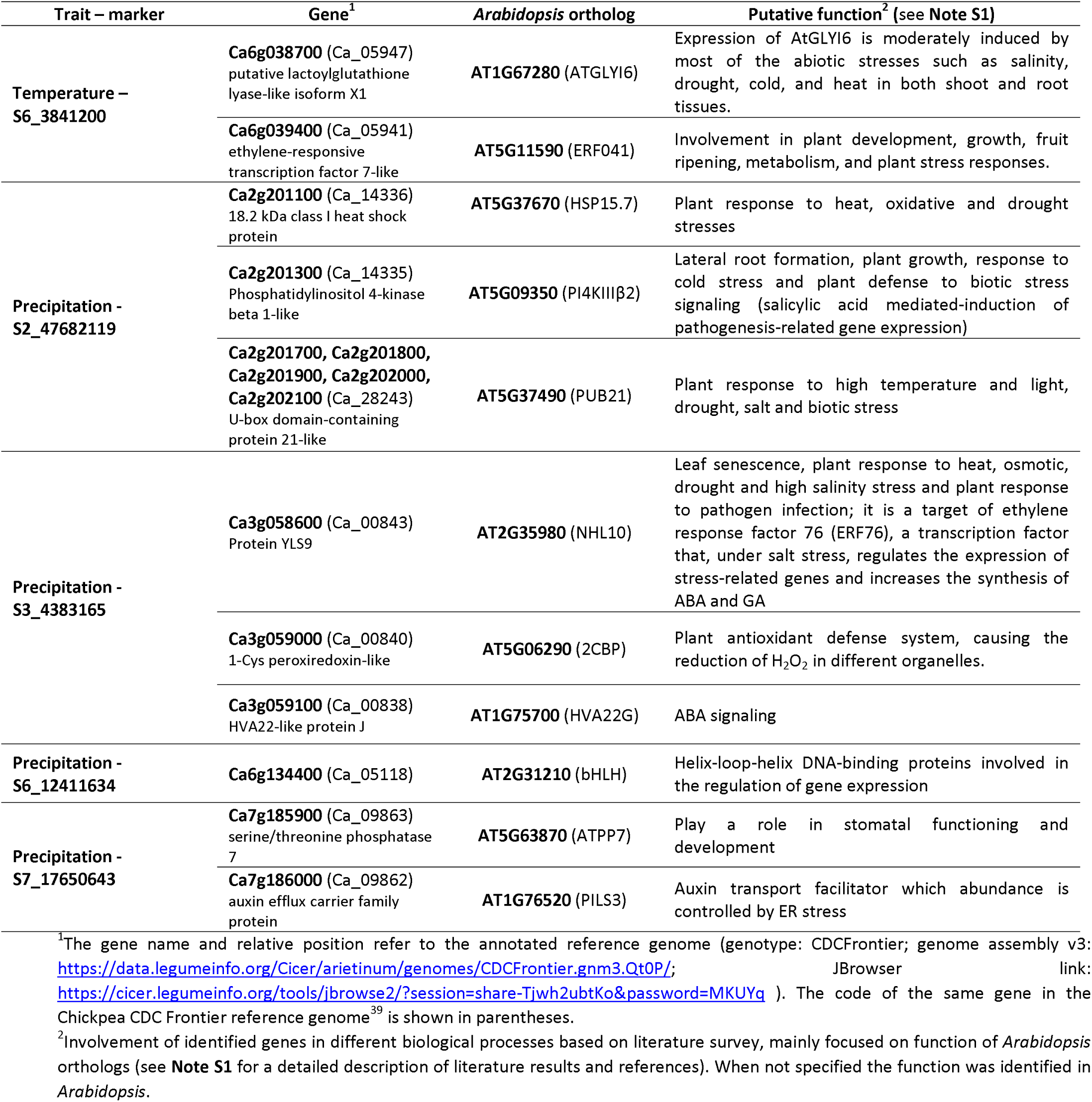
List of the most promising candidate genes for adaptation based on the function of orthologous genes in *Arabidopsis thaliana* and/or other species.

To visualize the geographic distributions of candidate adaptive alleles for heat and drought tolerance during the chickpea growing period we conducted an enriched RDA using based on the markers identified for temperature (94 loci) and precipitation (97 loci) variables using Storey FDR (q-value; q < 0.05) correction. The resulting RDA biplots delineate the adaptive genetic landscape, illustrating the correlations between genetic markers and the bioclimatic variables, as well as the correlation among the bioclimatic variables themselves (**Fig. 6A-B**). In the case of temperature variables, the enriched RDA (**Fig. 6A**) was significant (P<0.01), accounting for 19% of total variation, with RDA1 and RDA2 explaining 55.2% and 44.8% respectively of the total variation. The biplot highlights the RDA1 as positively correlated with the temperature in the drier quarter (bio9) and negatively correlated with the temperature in the wettest quarter (bio8). On the contrary RDA2 was positively correlated with both temperature variables. For precipitation variables, the enriched RDA (**Fig. 6B**) was significant (P<0.01), accounting for 20% of total variation, with RDA1 and RDA2 explaining 61% and 30.2% respectively of the total variation. RDA1 was negatively correlated with all precipitation variable considered, bio15 (precipitation seasonality), bio18 (precipitation in the driest quarter) and bio19 (precipitation in the coldest quarter), whereas RDA2 was negatively correlated with bio18 and positively correlated with bio19 and bio15.

**Fig. 6.**
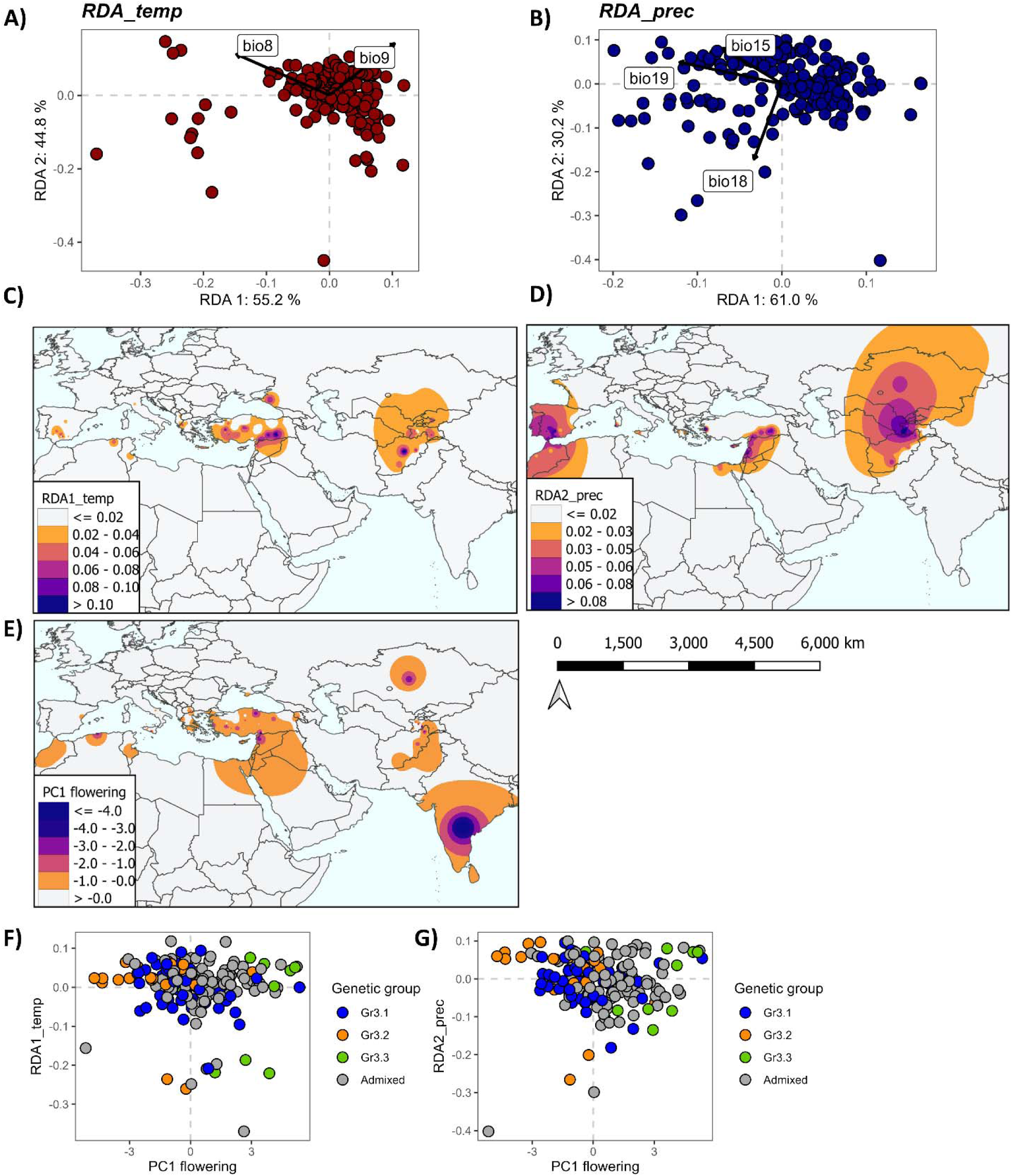
Chickpea adaptive landscape. **A,B)** enriched redundancy analysis (RDA) derived from adaptive loci (FDR q<0.05). Biplots showing genotype distribution in the adaptive space. **A)** RDA for temperature variable (*RDA_temp*) and **B)** RDA for precipitation variable (*RDA_prec*). **C,D,E)** Spatial distribution of *RDA1_temp*, *RDA2_prec* and PC1 of flowering time **F,G)** relation between PC1 flowering with *RDA1_temp* and *RDA2_prec*. Genotypes are colored in blue, orange and green to highlight the specific genetic groups: Gr3.1, Gr3.2, Gr3.3 (qi values >0.90).

Considering that most of our chickpea genotypes are sampled from latitudes between 30° and 45°, where chickpeas is cultivated as a spring crop, we decided to focus the analysis on critical variables such as bio9 and bio18, which at this latitude range represent the temperature and precipitation during the spring-summer period, respectively. Thus, we investigated the spatial distribution of the adaptive components such as RDA1 derived from the temperature variables (here named as *RDA1_temp*) and the RDA2 derived from precipitation variables (here named as *RDA2_prec*) (**Fig. 6 C-D-E**). Positive values of the *RDA1_temp* highlight the distribution of candidate adaptive alleles associated to heat stress (**Fig. 6C**). The spatial distribution of this component emphasizes the areas of Central Asia, particularly in the countries of Afghanistan and Tajikistan, as well as the southern part of Turkey. On the other hand, positive values of *RDA2_prec* highlight alleles potentially associated to low precipitation during the growing period (**Fig. 6D**). The spatial distribution of this component highlights the areas of Central Asia, as well as the southern east and southern west sides of the Mediterranean.

Considering the importance of flowering time as paramount adaptive trait for crops ^4^, we compared the spatial distributions of the adaptive alleles with the spatial distribution of flowering time, specifically using negative values for PC1, based on existing flowering data ^26^. The geographic areas of Central Asia, Southern Turkey and the Levant, resulted as overlapping regions where early flowering is used as adaptive strategy to escape heat and drought stress (**Fig. 6E**). Additionally, we identify the southwestern Mediterranean area, particularly in Morocco, where potentially adapted allele for drought and early flowering genotypes co-occurs. To further investigate the relationship between drought adaptation and flowering phenology, we compared the PC1 flowering scores with the adaptive RDA1_temp (**Fig. 6F**) and RDA2_prec components (**Fig. 6G**). Most of the early flowering genotypes, growing in latitudes between 30° and 45° (similar day length), were characterized by putatively heat and drought adaptive alleles (Chi square P<0.05). Moreover, early-flowering genotypes showing high *RDA1_temp* and *RDA2_prec* values belong mainly to Gr3.2 genetic group. These findings indicate that early flowering phenology characterized the adaptation strategy of this genetic group; by initiating the reproductive phase earlier, these genotypes complete their life cycle and reproduce before the onset of severe drought, increasing the likelihood of survival and successful reproduction in drought-prone environments. The same results are also found at K=6 partitioning (**Fig. S12**).

In conclusion, the application of landscape genomics and RDA enabled the characterization of genetic architecture of adaptive traits. By comparing the spatial distribution of adaptive alleles with the spatial variation of key agronomic traits, such as flowering time, this approach allowed to trace the adaptive strategies deployed along the main post-domestication diversification routes of chickpea. The identified GEA loci constitute a relevant reference set for interpreting the functional significance of agronomic QTLs identified across environmentally heterogeneous production systems, thereby expanding our understanding of the agronomic potential embedded in chickpea adaptive diversity. Furthermore, the landscape genomic framework demonstrated here highlights the value of exploiting georeferenced GenBank collections as a discovery platform for climatically relevant alleles, ultimately supporting more informed decisions in germplasm conservation and climate-adapted variety development.

## Materials and Methods

### Plant material

We assembled a panel of 532 chickpea lines obtained by single-seed descent (SSD) from diverse accessions ^25,26^, including 477 lines from the European and Mediterranean Chickpea Association Panel (EMCAP)^26^, most of which (392) were also part of the Training Core (T-CORE) in the INCREASE project ^4,25^. We combined the 477 EMCAP lines with the remaining 55 non-redundant lines from the INCREASE T-CORE. The final panel of 532 lines is listed with passport information in **Table S1.** Based on country of origin, we also grouped the lines into 15 geographic areas mostly in the Mediterranean basin, Middle East, and Central Asia. Latitude and longitude data for 208 lines were available from gene bank databases or collection sites (**Table S1**).

### Sequencing, variant calling and annotation

Genomic DNA was extracted from young leaves of single greenhouse-grown plants representing each line using the DNeasy Plant kit (Qiagen, Hilden, Germany). DNA libraries were prepared using the Illumina (San Diego, CA, USA) DNA PCR-free kit with > 350 ng of starting material. Equal volumes were pooled into final libraries that were quantified using a Qubit ssDNA assay kit (Thermo Fisher Scientific, Waltham, MA, USA) and then by real-time PCR against a standard curve using the KAPA Library Quantification Kit (Kapa Biosystems/Roche, Basel, Switzerland). Libraries were sequenced on an Illumina NovaSeq 6000 platform with S1, S2 and S4 flow cells in 150PE mode, yielding 15–30 million fragments per sample. Fragments were trimmed for base quality and adapters using fastp v0.21.0 and mapped to the *Cicer arietinum* reference genome (genotype CDCFrontier; genome assembly v3 https://data.legumeinfo.org/Cicer/arietinum/genomes/CDCFrontier.gnm3.QT0P/; Jbrowser link https://cicer.legumeinfo.org/tools/jbrowse2/?session=local-7zvicFS8v), kindly provided by the Cook laboratory (University of California, Davis), using BWA-mem v0.7.17. Overlapping reads were clipped using BamUtil v1.4.14 and duplicates were marked using Picard v2.17.11 MarkDuplicates. The mean coverage was 10×. Variants were called using HaplotypeCaller v4.1.9.0. Single-nucleotide polymorphisms (SNPs) were filtered as follows: Quality by Depth (QD) < 2.0; RMS Mapping Quality (MQ) < 40.0; Fisher Strand (FS) > 60.0; Mapping Quality Rank Sum Test (MQRankSum) < –12.5; Read Pos Rank Sum < –8.0; Strand Odds Ratio (SOR) > 3.0. Indels were filtered using the following parameters: QD < 2.0; FS > 60.0; Read Pos Rank Sum < –20.0. This identified 6,406,472 variants in total. According to the type of analysis, variants were also filtered by using the VCFtools ^44^ commands –remove-indels –max-missing 0.9 –mac 3 –minDP 3 –maf 0.05, retaining only chromosomal and biallelic sites, applying a minor allele frequency > 0.05 and a maximum missing value of 0.9. This narrowed the list to 639,648 SNPs that were used for genetic diversity analysis.

### Demographic analysis

Prior to population structure and ordination analyses, SNP datasets were thinned using the --thin flag in VCFtools ^44^, applying a minimum inter-SNP distance of 250 kbp to reduce redundancy arising from linkage disequilibrium among physically proximate markers. This approach is consistent with previous population genomic studies in self-pollinated species such as common bean ^9^.

Population structure was inferred using ADMIXTURE v1.3.0 ^45^, running 10 independent replicates for each value of K ranging from 2 to 8. Based on the K = 3 solution, an unrooted phylogenetic tree was reconstructed using the neighbor-joining (NJ) algorithm implemented in MEGA v11 ^46^, retaining only individuals with a cluster membership coefficient (q*i*) > 0.9. Tree topology reliability was assessed by bootstrap analysis with 10,000 replicates, and the resulting tree was visualized using the Interactive Tree Of Life (iTOL) platform ^47^. Principal Component Analysis (PCA) was performed using TASSEL v5 ^48^. For the 208 geo-localized lines, the spatial distribution of cluster-specific q*i* values was visualized through Inverse Distance Weighted (IDW) interpolation implemented in QGIS ^49^.

Genome wide Linkage Disequilibrium (LD) decay was estimated using the software PopLDdecay (v3.41) ^50^ considering a maximum distance of 2500Kbp.

### Landscape genomics

The 208 genotyped lines with known geographic coordinates were used for landscape genetic analysis. Missing genotypes were imputed using BEAGLE v4.1 ^51^, and the SNP dataset was subsequently thinned using a 10 kbp sliding window to limit the influence of linkage disequilibrium.

Bioclimatic variables were extracted from WorldClim (https://www.worldclim.org/) at 30 arcsecond resolution using the getData function from the R package *raster (version 4.4.3)*. A total of 19 bioclimatic variables were obtained for each genotype based on its geographic coordinates (**Table S2**). Isolation by distance (IBD) was assessed using a Mantel test ^52,53^ implemented in the R package *vegan* Community Ecology Package (version 2.7-1), computing the correlation between pairwise Euclidean genetic and geographic distances.

Redundancy analysis (RDA) was performed to quantify the influence of environmental variables on genome-wide genetic variation ^14^. To investigate potential adaptation to contrasting climatic drivers, the 19 bioclimatic variables were partitioned into two groups: temperature-related (bio1–bio11) and precipitation-related (bio12–bio19). Within each group, collinearity was assessed and variables with a variance inflation factor (VIF) > 2.5 were removed iteratively. This procedure resulted in the retention of two temperature variables (bio8 and bio9) and three precipitation variables (bio15, bio18, and bio19).

Population structure was accounted for by including the first three principal components from the genome-wide PCA as covariates, as these were not collinear with one another and were preferred over cluster membership coefficients (q*i*) for this purpose. The number of PCs to retain was determined by visual inspection of the eigenvalue scree plot, selecting the inflection point. Geographic effects were incorporated using the latitude and longitude of the geo-referenced genotypes.

Finally partial RDA (pRDA) was applied to partition the independent contributions of environment, population structure, and geography to total genetic variation ^14^. Genotype–environment associations (GEAs) were analyzed separately for precipitation and temperature using pRDA. The environmental variables were used as explanatory variables accounting for population structure, with geographic factors as covariates. Adaptive markers were identified using the *R* function *rdadapt* to calculate associations based on their extremeness along a distribution of Mahalanoibis distances estimated between each marker and the center of the RDA space using the first two RDA components ^14^. The *rdadapt* function returned p-values and q-values. Bonferroni correction (p = 0.05/number of tests) and Storey FDR (q-value; q < 0.05) correction were applied in either temperature and precipitation associations. Loci associated with temperature and precipitation were used as response variables for additional enriched RDA with bioclimatic variables as explanatory variables. Genotypes with enriched RDA score values were then used to map the spatial distribution of enriched RDA1 and RDA2 using IDW in QGIS. An additional GEA analysis was performed using the envGWAS ^36^ approach through the rMVP r package.

To investigate potential candidate genes, we used a window of 50 Kbp on either side of each significant SNP. This approach allows us to consider potential regulatory genes (e.g., enhancers) that might not be identified by the leading SNP. The sequences of candidate genes were used to identify *Arabidopsis thaliana* orthologs in TAIR. We also identified genes (chromosomal position) in the earlier chickpea reference genome, CDC Frontier v1.0 ^39^. Enriched terms for the described genes were identified using agriGO v.2.0 ^54^ *Arabidopsis thaliana* (TAIR10) annotated genes were used as background. The following parameters were set: hypergeometric test, multiple hypothesis test adjustment according to the Hochberg FDR procedure at significance level < 0.05, and minimum number of mapping entries = 3

### Phenotypic characterization of chickpea accessions for flowering

The 477 genotypes from the EMCAP collection were phenotyped in replicated field trails over 3 years (2019, 2020 and 2021) at the Research Centre for Cereal and Industrial Crops (CREA-CI) experimental station in Osimo, Ancona, Italy (latitude 43° 45′ 04′′; longitude 13° 49′ 98′′; 41 m above sea level) using a randomized complete block design with three replicates. Each plot consisted of a row of 10 plants, 1 m in length. We recorded the following flowering traits: first flower (FirstF), defined as the number of days from sowing date until 10% of plants have one open flower (flower banner, standard petal, is visible); full flower (FullF), defined as the number of days from sowing date until 50% of plants have one open flower (flower banner, standard petal, is visible); and first pod (FirstP), defined as the number of days from sowing date until 10% of plants have pods (pods are visible without removing flower petals). We used available flowering data ^26^ to carry out PCA in JMP v16.0 (SAS Institute, Cary, NC, USA).

## Acknowledgements

This research was supported by funding from the European Union Horizon 2020 Research and Innovation program through the INCREASE project (Grant n. 862862), the MUR - PON R&I 2014-2020 through the project RESO (Grant n. ARS01_01224), and from the Polytechnic University of Marche (2021-2023).

## Author contributions

RP designed the research nd contributed to data analysis and interpretation; LR, EBi, EBe, LN, SP, CS and CB developed the SSD lines; CB collected leaf tissues; SP performed DNA extraction; LDA and EV conducted sequencing; AV performed primary bioinformatic analysis; LR performed data analysis; MRod, AP, Ebi, RB and AB contributed to data analysis and discussion of the results; EBi and GF carried out the functional survey of candidate genes; LR, GF and EBi wrote the Supporting Information; LR, GF, EBi and RP wrote the manuscript; MD, MRos and MRod a contributed to the writing and drafting. All authors read and approved the article.

## Data availability

The Whole Genome Sequencing (WGS) raw data were uploaded to NCBI database with accession number of PRJNA1070423. The bioinformatic pipelines used in this work are available at https://github.com/LorePlant/Landscape_chickpea.

