## Supplemental Figures for "Landscape genomics highlights the adaptive evolution of chickpea across the Silks Roads"

Rocchetti et al.

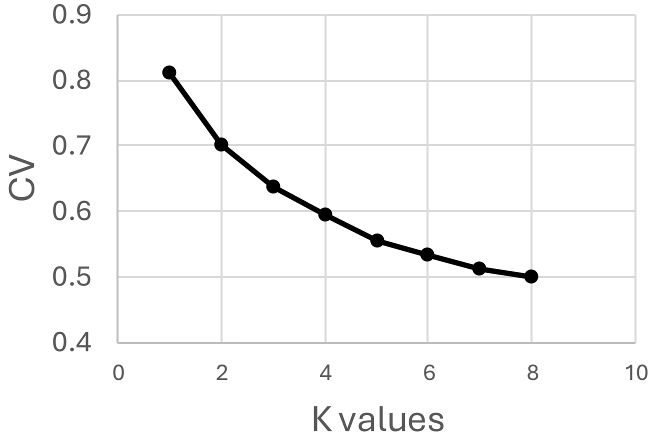


**Figure S1:** Cross-validation plot of admixture analysis. Admixture with cross-validation for K from 1 to 8.

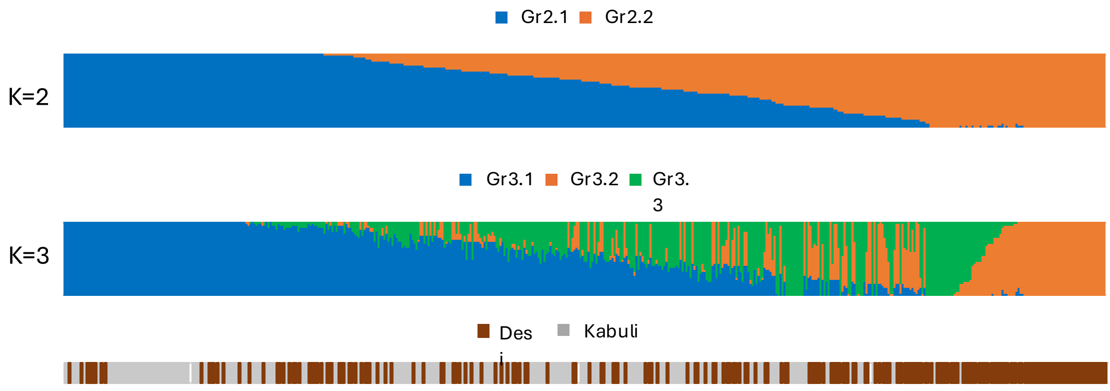


**Figure S2:** Population structure analysis run with ADMIXTURE based on 2,050 SNPs single nucleotide polymorphisms (SNPs) across 532 Chickpea accessions. Ancestry bar plot at K (2-3) and differentiation between desi and kabuli types. Across the three bar plots genotypes are sorted according to K=2 subdivision.

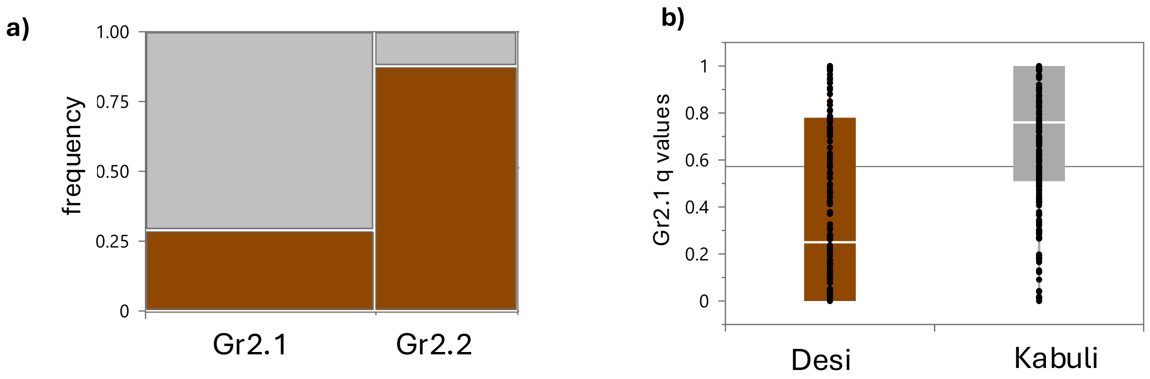


**Figure S3**: a) Mosaic plot of desi and kabuli frequency within Gr2.1 and Gr2.2 groups (*qi>0.9*). **b)** One-way Anova desi/kabuli types for Gr2.1 *qi* values

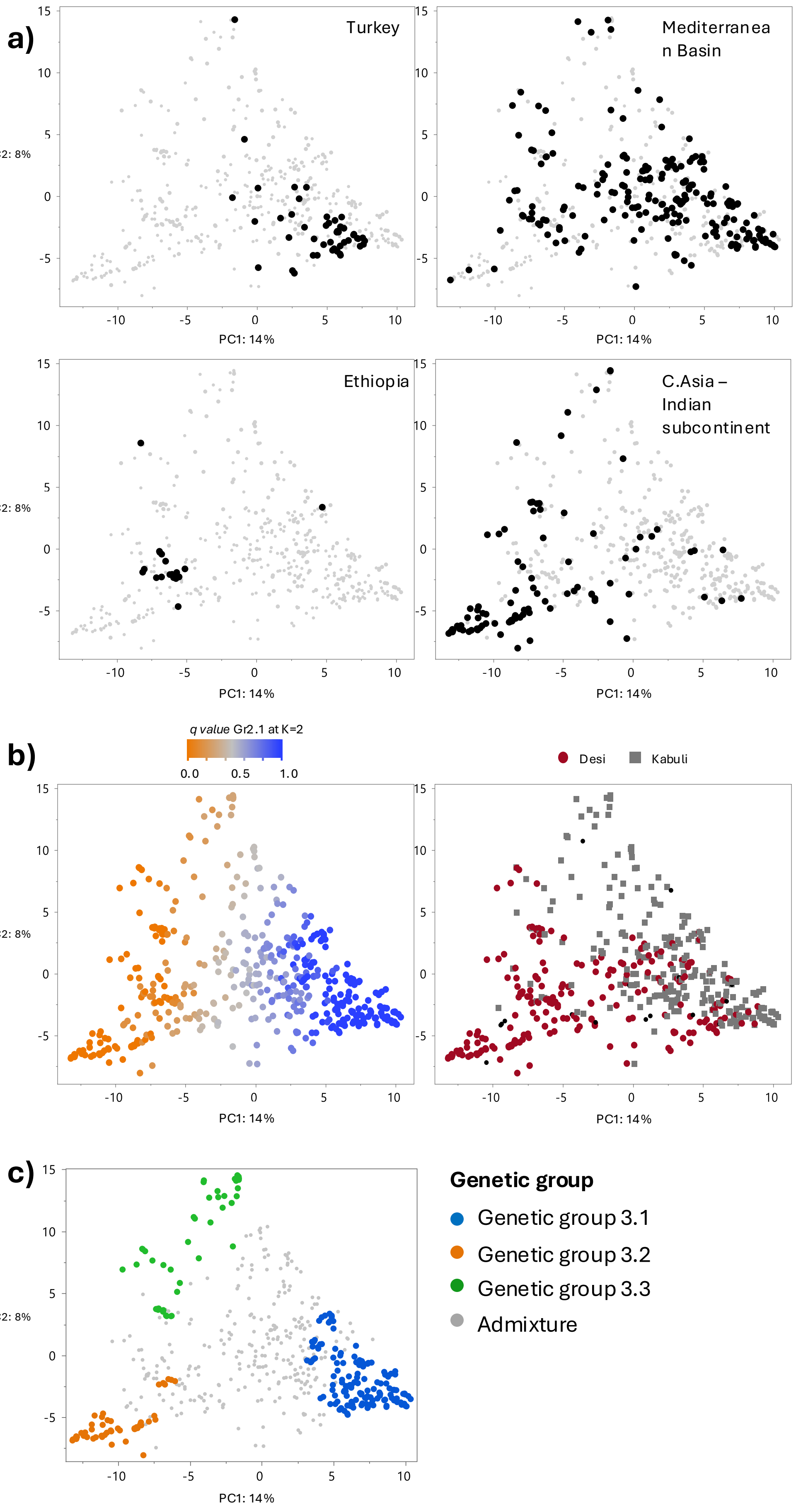


**Figure S4**: Principal component analysis with 2,050 SNPs single nucleotide across 532 genotypes. **a)** score plots highlighting four important geographic area of chickpea domestication history: Turkey, Mediterranean Basin, Central Asia together with Indian subcontinent and Ethiopia. **b)** score plots highlighting Gr2.1 and Gr2.2 differentiations and desi/kabuli dichotomy. **c)** representation of the pure *(qi>0.9)* Gr3.1, Gr3.2 and Gr3.3.

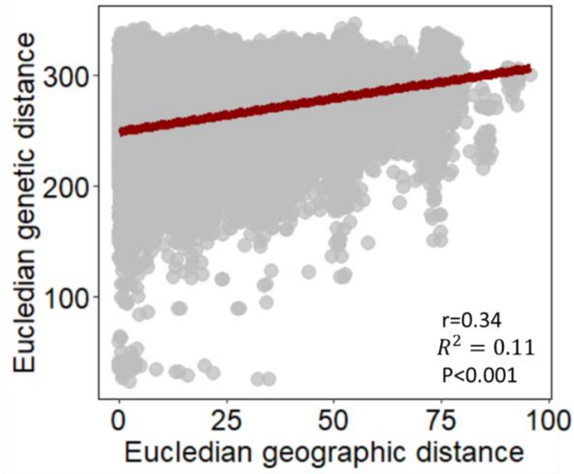


r=0.34

$R^2$

$= 0.11$

P<0.001

**Figure S5:** Mantel test between Euclidean genetic distance and Euclidean geographic distance.

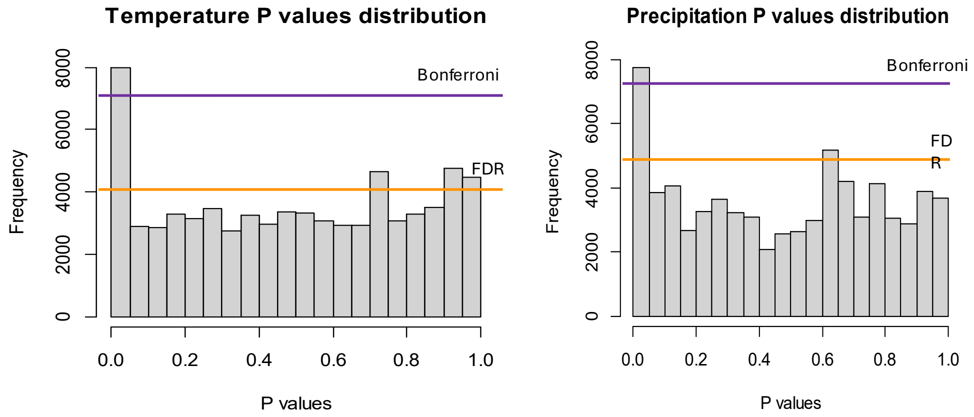


**Figure S6:** P values distributions for temperature and precipitation associations. In red Bonferroni threshold and in blue the FDR threshold.

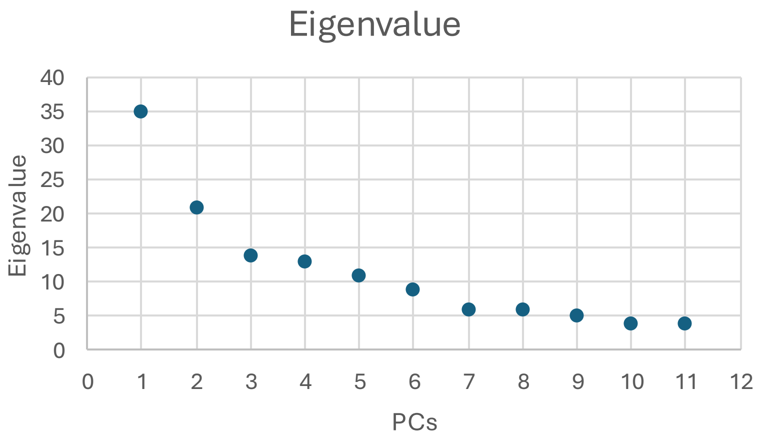


**Figure S7:** Eigenvalue decay derived from the PCA conducted with 2,052 SNPs

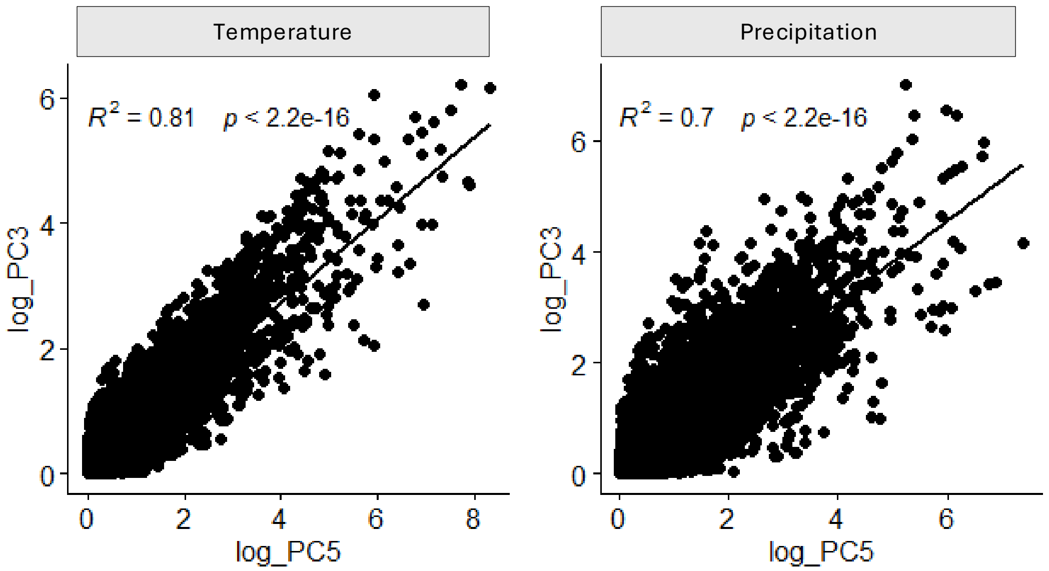


**Figure S8:** Comparison between GEAs identified for temperature and precipitation, while correcting for population structure using three PCs( log_PC3) and five PCs (log_PC5)

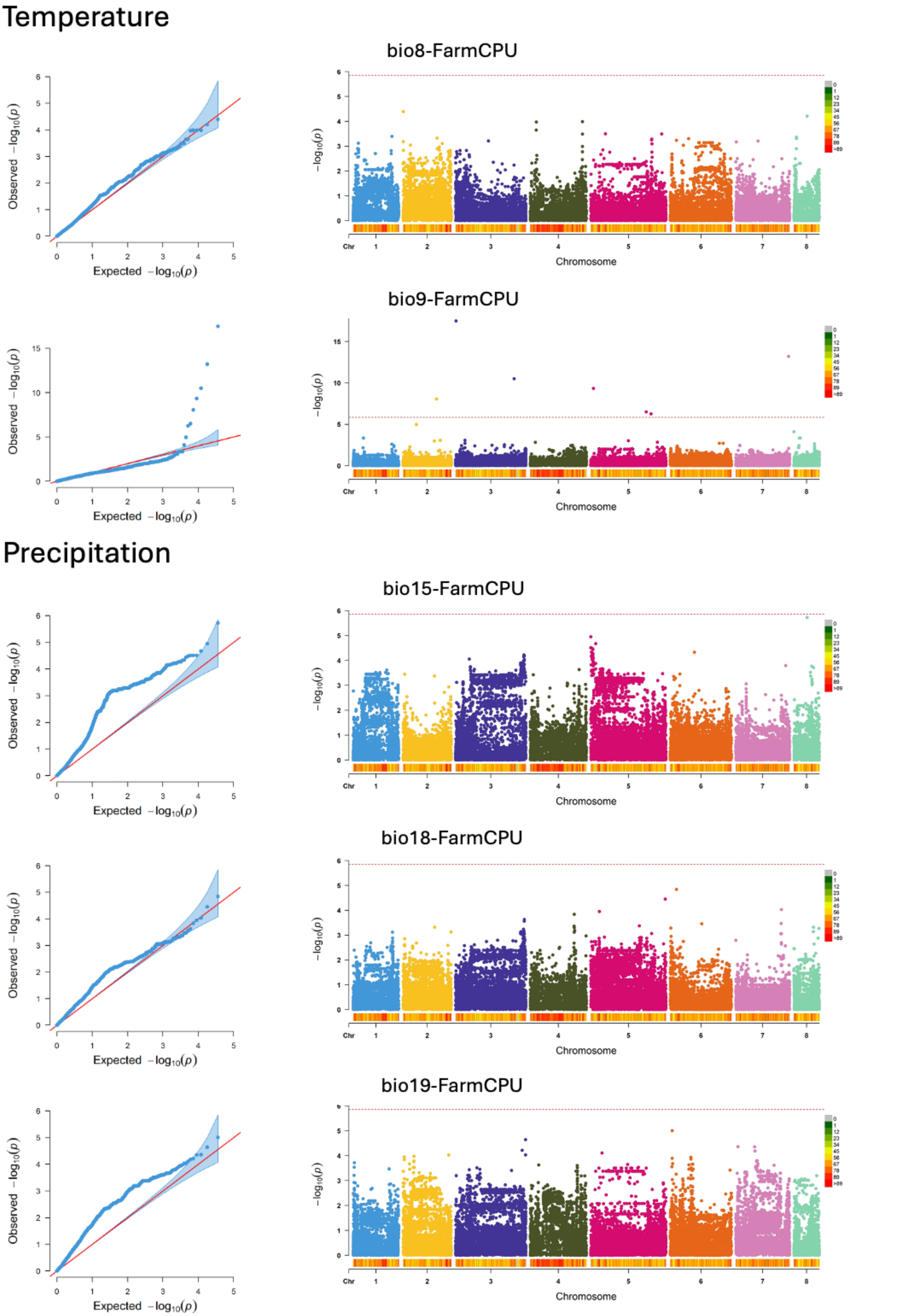


**Figure S9:** Genotype Environment Association (GEAs) found using envGWAS for temperature (bio8 and bio9) and precipitation (bio15, bio18,bio19) variables.

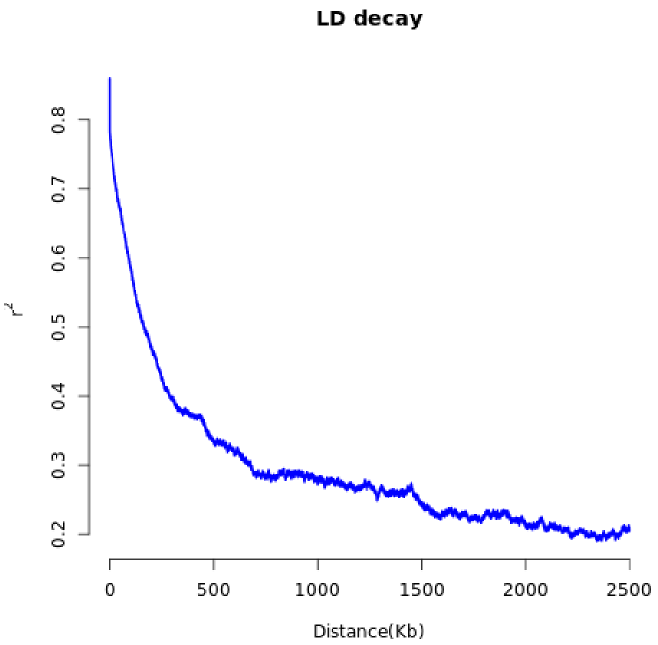


**Figure S10:** Linkage disequilibrium decay estimated genome wide, considering a maximum window of 2500Kbp

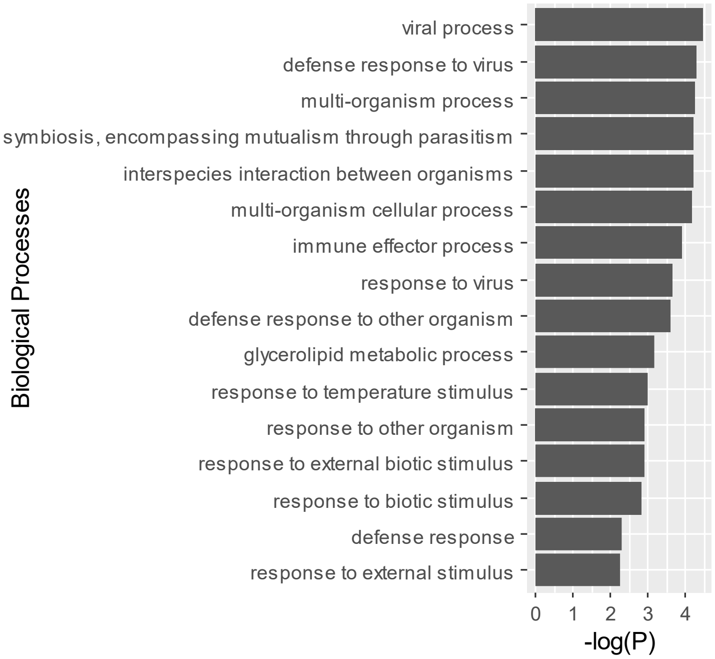


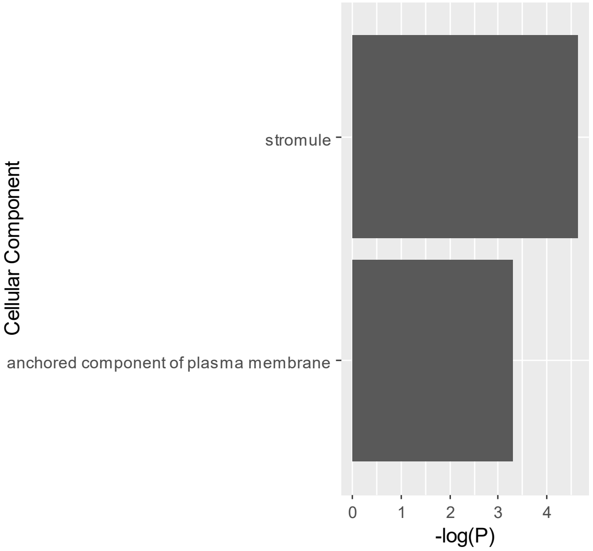


**Figure S11:** GO enrichment against *Arabidopsis thaliana* (TAIR10) genome.

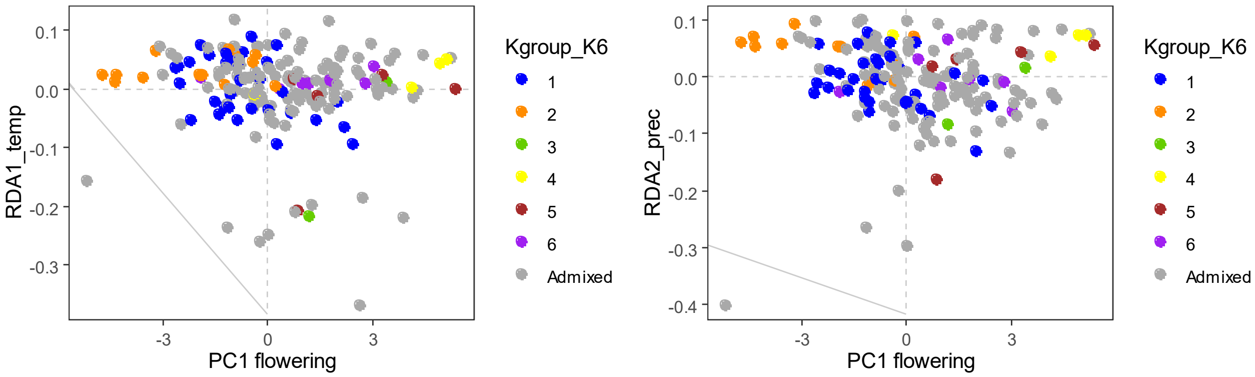


**Figure S12:** Relation between PC1 flowering with RDA1_temp and RDA2_prec. Genotypes are colored in blue, orange, green, yellow, brown and purple to highlight the specific genetic group Gr6.1, Gr6.2, Gr6.3, Gr6.4, Gr6.5 and Gr6.6 (qi values >0.90) respectively.
