## Supplementary material for "Landscape genomics highlights the adaptive evolution of chickpea across the Silks Roads": Note S1

**Notes S1 Functional annotation of candidate genes**

The functions of a set of candidate genes for adaptation were investigated based on the descriptions in the annotated reference genome (genotype: CDCFrontier; genome assembly v3: https://data.legumeinfo.org/Cicer/arietinum/genomes/CDCFrontier.gnm3.QT0P/; JBrowser link: https://cicer.legumeinfo.org/tools/jbrowse2/?session=share-Tjwh2ubtKo&password=MKUYq) and according to the function of the orthologous genes in *Arabidopsis thaliana* (TAIR database).

A list of 54 candidate genes was established by considering nine windows of 100 kb (50 kb upstream and downstream of SNP markers associated with temperature and precipitation variables) for all the identified SNPs.

*Temperature - S6_3841200*

Furtheen gense were fund in the 100 Kb window centered on SNP S6_3841200 associated with temperature variables.

Ca6g038600 encodes a putative cytoplasmic methyltransferase belonging to the S-adenosyl-L-methionine (SAM)- dependent methyltransferase family. Its A. thaliana ortholog, AT5G51130, is involved in protein amino acid methylation, displays diurnally rhythmic expression [1].

Ca6g038700 encodes a glyoxalase I / lactoylglutathione lyase involved in methylglyoxal detoxification. The A. thaliana ortholog AT1G67280 is involved in the catalysis of the first step of the glyoxal pathway. The function of this gene is associated with the response to cold, salinity and metal ion binding [2]

Ca6g038800 and Ca6g038900 are two close homologs that encodes for sugar transport proteins involved in monosaccharide transport and uptake during seedling growth and stress responses. AT1G77210 is the ortologs in A. thaliana, it encodes for a plasma-membran monosaccharide transporter. It belongs to the STP (Sugar Transport Protein) family and functions as a proton-coupled galactose transporter. The protein is part of the major facilitator superfamily (MFS) and is expressed in multiple tissues, suggesting a role in carbohydrate uptake and distribution [3].

Ca6g039100 encodes for the CRS2-associated factor 1. Its ortholog in A. thaliana (AT1G77210) also known as CAF1 encodes an RNA-binding protein containing a CRS1/YhbY (CRM) domain. It functions in the splicing of chloroplast group II introns, helping remove introns from specific chloroplast transcripts such as rpoC1 and clpP-1 [4].

Ca6g039200 probably encodes for glycosyltransferase isoform X4. The A. thaliana ortholog, AT5G19670, is annotated as a putative hydrolase-related protein.

Ca6g039300 encodes the α/β hydrolase superfamily. Its otholog in A. thaliana is AT2G44970 is involved in intracellular protein transport and hydrolitic activity that act on ester bonds.

Ca6g039400 encodes the ethylene-responsive trascription factpor 7 like. The A. thaliana ortholog is AT5G11590 and it is involved in stress-related and hormone-responsive pathways. It encodes an AP2/ERF family transcriptor factor, this domain has been shown in various proteins to be necessary and sufficient to bind the GCC-box that is essential to the response to ethylene [5,6].

Ca6g039500 has no informtion but its ortholog in A. thaliana (AT1G40087) encodes for transposase proteins that are necessary the an efficent DNA transposition.

No information is available for the Ca6g039500. Nevertheless, the AT3G20980 encodes a Gag-Pol-related retrotransposon family protein in A. thaliana.

Ca6g039700 putatively encodes a protein phosphatase 1 regulatory subunit 7-like isoform X1. The A. thaliana ortholog, AT5G19680, codes for a leucine reach repeat (LRR) protein family. This protein domain appears to provide a structural framework for the formation of protein-protein interactions [7]. Proteins characterized by LRRs are involved in different biological processes, including signal transduction, cell adhesion, DNA repair, recombination, transcription, RNA processing, disease resistance, apoptosis, and the immune response [8].

Ca6g039800 and its ortholog AT1G77170 (A. thaliana) putatively enconde a pentatricopeptide repeat (PPR) superfamily protein [9]. These PPR proteins typically localize to mitochondria and are involved in RNA processing and organellar gene expression in mitochondria or chloroplasts.

No information is available for Ca6g039900.

*Precipitation - S2_47682119*

Thirteen genes were found in the 100-kb window centered on SNP S2_47682119 associated with precipitation variables. Three genes (Ca2g201200, Ca2g201400 and Ca2g202200) are uncharacterized.

Ca2g201100 encodes a class I 18.2-kDa heat shock protein. The AT5G37670 (HSP15.7) gene is the ortholog in *A. thaliana*. It encodes an HSP20-like chaperone superfamily protein involved in protein complex oligomerization, protein folding, and cellular responses to hypoxia, heat, $H_2O_2$, reactive oxygen species and salt stress. Sewelam et al. (2019) analyzed the expression of different HSPs in *A. thaliana* ecotype Columbia (Col-0, wild type) under different stress treatments (heat, drought, salinity and oxidative stresses imposed by $H_2O_2$) and found that the HSP15.7 gene was induced by heat and oxidative stress. HSP15.7 was also detected by Charng et al. (2007) as a heat stress responsive gene in Arabidopsis. Similarly,Rizhsky et al. (2004) found that HSP15.7 was among the transcripts present at high levels in leaves of Arabidopsis plants subjected to a combination of drought and heat stress.

Five closely linked genes (Ca2g201700, Ca2g201800, Ca2g201900, Ca2g202000 and Ca2g202100) encode homologs of U-box domain-containing protein 21. The ortholog gene in Arabidopsis, AT5G37490, encodes a plant U-box type E3 ubiquitin ligase (AtPUB21, AtCMPG5). Fei et al. (2019) carried out transcriptome profiling in Arabidopsis roots and compared the differences in gene expression at lower, higher and normal plant growth temperatures to acquire new knowledge on how root growth adapts to elevated ambient temperatures. They found that AtPUB21 was among the genes dominantly downregulated in response to high temperature. AtPUB21 was also among the genes upregulated in cell suspension cultures after 30 mins of strong illumination [14]. Xu & Huang (2017) analysed the leaf transcriptome of creeping bentgrass (*Agrostis stolonifera*) wildtype cultivar Penncross and a mutant transformed with the isopentenyl transferase (*ipt*) gene (line S41, known to overexpress cytokinin and positively regulate drought tolerance). The AtPUB21 ortholog was among the downstream genes showing significant changes of expression levels between S41 and wild-type under drought stress, indicating that the improved drought tolerance in *ipt*-transgenic plants was related to ubiquitin-mediated proteolysis cascades. Numerous other examples show that U-box E3 genes are involved in responses to abiotic stress, such as drought, light and salt (Cho et al., 2008; Bergler & Hoth, 2011; Seo et al., 2012; Sharma et al., 2013.; Woodson et al., 2015; Adler et al., 2017; Song et al., 2017; Wang et al., 2020). In Arabidopsis, U-box type E3 ubiquitin ligase (PUB21) is involved in biotic stress responses [24]. Orthologous genes in tobacco (NtCMPG1) and tomato (Cmpg1) are essential for plant defence and disease resistance [25]. The orthologous gene in rice (Os06g0248500) was differentially expressed in the blight-resistant line *bbr1* [26].

Ca2g201300 encodes a phosphatidylinositol 4-kinase β1-like protein. The Arabidopsis ortholog AT5G09350 encodes a phosphatidylinositol 4-OH kinase (PI4KIIIβ2) involved in phosphatidylinositol biosynthesis, pollen tube growth and root hair cell tip growth. Rubilar-Hernández et al. (2019) showed that the gene is involved in lateral root formation. Loss-of-function of PI4KIIIβ1 (AT5G64070) and PI4KIIIβ2 triggered lateral root primordium formation and endocytic trafficking toward the vacuole, indicating that these genes negatively regulate the formation of lateral roots. This function is interesting in the context of abiotic stress given that an increase in the contact area between the roots and rhizosphere can improve the uptake of water and nutrients from the soil. Antignani et al. (2015) showed that PI4KIIIβ2 is also involved in plant defence signaling in Arabidopsis. Along with PLANT U-BOX13 (PUB13) and PI4KIIIβ1, it is a negative regulator of salicylic acid-mediated induction of pathogenesis-related gene expression. The double mutant for phosphatidylinositol-4-kinases (pi4kIIIβ1β2) was also shown to accumulate salicylic acid in upper tissues leading to the dwarf phenotype and indicating a role in plant growth [29]. Delage et al. (2012) showed that PI4KIIIβ1 and PI4KIIIβ2 have a crucial role during cold stress: along with PI4KIIIα1, they provide substrates to phosphoinositide-dependent phospholipase C (PI-PLC) during cold stress. PI-PLC produces diacylglycerol (DAG) which can be phosphorylated to phosphatidic acid (PtdOH). The latter is significantly depleted in the pi4kIIIβ1β2 double mutant exposed to cold stress [30].

Ca2g201500 encodes an A49-like RNA polymerase I associated factor. The Arabidopsis ortholog AT3G13940 encodes a DNA binding/DNA-directed RNA polymerase involved in RNA polymerase I preinitiation complex assembly and transcriptional elongation by RNA polymerase I. It was among the most significant 50 genes expressed differentially in shoots and roots of cesium-intoxicated compared to non-intoxicated Arabidopsis plants [31]. This gene was differentially expressed in both single and double copper-iron deficiency compared to control conditions in Arabidopsis [32].

Ca2g201600 encodes a cysteine-rich receptor-like protein kinase 25. Its orthologous gene in Arabidopsis (AT5G37660) encodes plasmodesmata-located protein 7 (PDLP7) involved in plant immunity against bacteria [33]. Palomar et al. (2021) showed that PDLP7 is also involved in the germination and post-germination growth of Arabidopsis seeds under salinity stress.

Ca2g202300 encodes a putative membrane lipoprotein. The orthologous gene in Arabidopsis (ZENGDA SMALL PEPTIDE 1, ZSP1) encodes a plant-specific small peptide involved in the regulation of organ size [35]. Loss-of-function mutant *atzsp1-1* exhibited small organs, whereas AtZSP1 overexpression plants (mutant p35S:AtZSP1#1) produced larger organs. Differentially expressed genes in the shoots of both these mutants were enriched in the cytokinin pathway, suggesting that AtZSP1 affects shoot size by changing cytokinin levels [35]. The same study also indicated that AtZSP1 interacts with REPRESSOR OF CYTOKININ DEFICIENCY 1 (ROCK1) to regulate cytokinin levels, and thus organ size.

*Precipitation - S3_4383165*

Fifteen genes were found in the 100-kb window centered on SNP S3_4383165 associated with precipitation variables. Four of the genes (Ca3g058400, Ca3g058900, Ca3g059200 and Ca3g059700) are uncharacterized.

Four genes within this window (Ca3g058300, Ca3g058500, Ca3g058600, and Ca3g058700) encode YELLOW-LEAF-SPECIFIC GENE 9 (YLS9)-like proteins, belonging to the LEA hydroxyproline-rich glycoprotein family. Ca3g058500 and Ca3g058700 are co-orthologs of Arabidopsis AT3G11650, whereas the orthologs of Ca3g058300 and Ca3g058600 are AT3G11660 and AT2G35980, respectively. AT3G11660, AT3G11650 and AT2G35980 are paralogous Arabidopsis genes encoding proteins related to tobacco hairpin-induced gene (HIN1) and Arabidopsis non-race specific disease resistance gene (NDR1), known as NDR1/HIN1-LIKE proteins or NHL genes. AT3G11660, AT3G11650 and AT2G35980 are NHL1, NHL2 and NHL10, respectively. Both HIN1 in tobacco and NDR1 in Arabidopsis are known to be involved in responses to pathogen infection [36,37].

NHL10 expression is associated with leaf senescence and the avirulent Cucumber mosaic virus (CMV)-induced hypersensitive response (HR), and are induced by exogenously applied spermine (Spm), which also induces pathogenesis-related genes in tobacco [38]. A role for NHL1 and NHL2 in response to bacterial pathogens has been suggested [39]. Maldonado et al. (2014) overexpressed four members of the NHL gene family previously studied in Arabidopsis (NHL1, NHL2, NHL8 and NHL25) in soybean (*Glycine max*) roots and investigated the effect on resistance to the soybean cyst nematode (*Heterodera glycines*). They found that NHL1 and NHL8 reduced the soybean cyst nematode population by ~50% and the overexpression of NHL1 and NHL8 induced the expression of marker genes for jasmonic acid and ethylene [40].

NHL genes are also involved in abiotic stress responses (Lee et al., 2006; Bao et al., 2016; Liu et al., 2020; J. Wang et al., 2022; Zhang et al., 2022) Chickpea NHL10 (Ca3g058600, named Ca_00843 in the chickpea CDC Frontier reference genome; Varshney et al., 2013) was among the differentially expressed proteins identified between control and drought stressed root tissues of the JG 11+ genotype ([47]). The poplar genes Potri.006G204300 and Potri.016G071600 (orthologs of NHL10) are involved in the response to salinity stress, and are among the targets of ethylene response factor 76 (ERF76), a transcription factor that upregulates the expression of stress-related genes and increases the ability of the plant to synthesize ABA and gibberellin, which results in stronger tolerance to salt stress [48]. The soybean NHL10 ortholog GmSYP24 is strongly and rapidly induced by drought stress, and its overexpression in soybean or heterologous expression in Arabidopsis confers insensitivity to osmotic/drought stress and high salinity [49]. Shahbaz et al. (2023) investigated the effects of environmental heat stress on NHL gene expression in maize and found that the ZmNHL10 gene is significantly upregulated under heat stress in genotypes B-73 and W-22.

The Ca3g058800 gene encodes a eukaryotic aspartyl protease. The Arabidopsis ortholog AT3G52500 is involved in numerous biological processes (cell division, cell wall organization or biogenesis, flavonoid biosynthesis, growth, oxoacid metabolism, and response to bacteria, jasmonic acid, light and wounding). Raggi et al. (2015) analyzed the transcriptome of rosette leaves in two independent Arabidopsis 35S:AnPGII transgenic lines expressing a mutated version of the *Aspergillus niger* polygalacturonase II with reduced activity [52]. The pectin composition was altered in the transgenic plants, resulting in severe growth defects. The AT3G52500 transcript levels decreased significantly in the rosette leaves of both transgenic lines compared with the wild type, indicating a role in plant response to cell wall damage.

The Ca3g059000 gene encodes a 1-Cys peroxiredoxin-like protein. The Arabidopsis ortholog (AT5G06290, 2-CysPrxB, 2CPB) is a 2-Cys peroxiredoxin (2-CysPrxB) that contains two catalytic cysteine residues. It is involved in cell redox homeostasis, response to cold and oxidative stress, and $H_2O_2$ catabolism. Peroxiredoxins have been described as key components of the plant antioxidant defence system that reduce $H_2O_2$ levels in chloroplasts [53]. In Arabidopsis, At2-CysPrxB (At5g06290) maintains the water-water cycle for proper $H_2O_2$ scavenging, and double mutants deficient in 2-CysPrxA and 2-CysPrxB accumulated reactive oxygen species resulting in the photo-bleaching of leaves during high light stress [54]. Mao et al. (2018) investigated the function of the 2-CysPrxB ortholog in rice (OsPRX2) particularly its effect on potassium-deficiency tolerance. The overexpression of OsPRX2 led to stomatal closure and higher tolerance, whereas the knockout of OsPRX2 caused stomatal opening during potassium deficiency and defects in leaves [55]. Gallardo-Martínez et al. (2023) identified 2-cys peroxiredoxins A and B as two members of the plastid redox system involved in early plant development, revealing that the Arabidopsis double mutant for these genes is deficient for seed development and embryogenesis.

Ca3g059100 encodes HVA22-like protein J. The Arabidopsis ortholog AT1G75700 acts upstream of or directly in the cell cycle, protein complex assembly, sulfur compound biosynthesis and the responses to cold and light. Although the function of the HVA22-like family is unclear, its members are induced by ABA in barley and Arabidopsis [57,58]. ABA signaling induces drought and osmotic stress tolerance [59] and HVA22-like genes are involved in responses to abiotic stress such as salinity, drought and cold. Arabidopsis ATHVA22 is strongly induced under drought conditions and helps to prevent desiccation (C. N. Chen et al., 2002; Amara et al., 2022). HVA22-like genes in barley are also involved in abiotic stress responses [61]. Lyu et al. (2021) showed that HVA22 was upregulated in faba bean under cold stress. In cotton, GhHVA22E1D responds to salt and drought stress [63]. Wai et al. (2022) confirmed that the SlHVA22 gene family is involved in abiotic stress tolerance in tomato.

The Ca3g059300 gene encodes an α/β-hydrolase family protein. Its Arabidopsis ortholog is AT3G11620, which facilitates the intracellular distribution of lipid droplets (TAIR DATABASE).

The Ca3g059400 and Ca3g059500 genes encode two ovate family proteins. The Arabidopsis orthologs AT3G52525 and AT5G04820 encode ovate family protein 6 (OFP6) and 13 (OFP13), respectively, which are transcriptional repressors. Ovate family proteins have been shown to regulate multiple aspects of plant growth and development [65] in Arabidopsis [66], tomato [67], rice [68], potato [69], pepper [70] and radish [71]. Ma et al. (2017)showed that OsOFP6 is also involved in drought and cold stress responses. OsOFP6 overexpression increased cold and drought stress tolerance, whereas RNAi knockdown conferred greater sensitivity [72].

The Ca3g059600 gene encodes a CCR4-associated factor 1 homolog 11-like protein. The Arabidopsis ortholog AT3G44260 is related to yeast CCR4-associated factor 1 (CAF1a), which regulates mRNA deadenylation and responses to pathogens [73]and wounding [74]. CAF1a expression is also an early and transient response to cold stress [75].

*Precipitation - S6_12411634*

Five genes were found in the 100-kb window centered on SNP S6_12411634. This marker is associated with precipitation variables. The gene Ca6g134200 and Ca6g134500 are poorly characterize.

Ca6g134100 and its orthologous gene in Arabidopsis thaliana (AT4G37340) encode the Cytochrome P450 superfamily protein. Cytochrome P450 enzymes are a superfamily of haem-containing mono-oxygenases that are found in all kingdoms of life. In plants these proteins are involved in the biosynthesis of several compounds such as defensive compounds through the monooxygenation of a carbon atom [76] .

Ca6g134200 is poorly characterized; however, its Arabidopsis thaliana orthologs (AT2G25670) are putatively involved in copper ion binding.

Ca6g134300 encodes a homeobox-leucine zipper family protein belonging to the HD-ZIP IV family. AT3G03260 is the orthologous gene in A. thaliana and it is involved in the regulation of epidermal differentiation and it plays a role in response to biotic and abiotic stresses [77].

Ca6g134400 encodes the basic helix-loop-helix (bHLH) DNA-binding superfamily protein. The same domain is encoded by AT2G31210 in A. thaliana, and it is represented by sequence specific DNA-binding proteins that have the role of transcription factors and are involved in the regulation of gene expression. As reported by Heim et al. (2003), the bHLH superfamily protein is highly diversified, suggesting that the proteins belonging to this domain regulate a wide range of developmental processes and stress responses in plants.

Ca6g134500 is poorly characterized. Its orthologs gene A. thaliana is AT3G47630 (Tam41). It is a mitochondrial phosphatidate cytidylyltransferase. It is involved in the stabilization of the supercomplexes of the mitochondrial respiratory chain in the inner membrane of the organelle and as shown by Leviatan et al. (2013) it is involved in the cold stress response.

*Precipitation - S7_17650643*

Within a 100kb window surrounding the SNP S7_17650643, seven genes associated with precipitation variables were identified.

Ca7g185900 encodes the serine/threonine phosphatase 7. The A. thaliana orthologs is AT5G63870 and it encodes a nuclear localized serine/threonine phosphatase (PP7) that appears to be regulated by redox activity and is a positive regulator of cryptochrome mediated blue light signalling [80]. In addition, as propose by Genoud et al. (2008), PP7 also modulates phytochrome-dependent light responses likely through interactions with light-signaling components such as NDPK2.

Ca7g186000 and its orthologous gene in A. thaliana (AT1G76520) encode an auxin efflux carrier family protein (PILS3). These kinds of carriers are auxin specific and are localized at the basal end of the auxin transport-competent cell [82]. The PILS3 is likely involved in the regulation of plant growth and development since it contributes to auxin distribution within tissues[83].

Ca7g186100 and Ca7g186200 putatively encode a diacylglycerol O-acyltransferase 2-like. The gene AT3G51520 encodes the DGAT2 in A. thaliana, an enzyme catalyzing the final step in triacylglycerol biosynthesis. In addition, DGAT2 seems involved in the biosynthesis of TAGs, a key storage lipid [84].

Ca7g186300 is poorly characterize. The A. thaliana orthologous gene AT4G10520 encodes a subtilase family protein, which in plants are involved in protein maturation, turnover, and signaling processes.

Although this specific gene has not been experimentally characterized, it is predicted to exhibit serine-type protease activity and participate in proteolytic pathways relevant to plant development and stress responses [85].

Ca7g186400 putatively encodes a kiesi-4-like isofom X2. AT5G65930 (ZWI) encodes a kinesin-like calmodulin-binding motor protein in Arabidopsis thaliana. It functions in microtubule-based processes and is required for normal trichome branching and pollen germination [86].

Ca7g186500 is poorly characterized. The A. thaliana orthologous gene AT1G78690 encodes a phospholipid/glycerol acyltransferase family protein predicted to function as a lysoglycerophospholipid O-acyltransferase involved in glycerophospholipid metabolism and membrane lipid remodeling [87].
